# A Translocation within the *Ogataea* Species Complex Alters Local Subtelomeric Chromatin while Maintaining Overall Genome Organization

**DOI:** 10.64898/2026.03.05.709876

**Authors:** Tiffany J. Lundberg, Nickolas M. Lande, Daniel Tourevski, Riley Figueroa, Sara J. Hanson, Andrew D. Klocko

## Abstract

Eukaryotic genomic DNA is packaged in the nucleus as chromatin – a DNA-protein aggregate regulating genome function, including transcription. Chromatin is classified as either active euchromatin or silent heterochromatin, with each marked by distinct histone post-translational modifications (PTMs). Chromatin composition also mediates genome organization, including how heterochromatin aggregates at the nuclear periphery while euchromatin localizes to the nucleus center. In fungi, heterochromatic loci cluster, including independent centromere and telomere clusters that form the Rabl chromosome conformation. However, it is unknown if chromatin composition and genome organization are conserved in closely related fungi, and how these features are impacted by large-scale chromosomal rearrangements. Here, we examined differences in histone PTM deposition, gene expression, and genome organization in two yeast species from the order Pichiales, which diverged from the common ancestor shared with *Saccharomyces cerevisiae* more than 200 million years ago. We focused on *Ogataea polymorpha*, which is used for industrial protein production, and *Ogataea haglerorum*, an isolate of which harbors a translocation between chromosomes 1 and 6. We show that the enrichment of three activating PTMs – the trimethylation of lysine 4 of histone H3 (H3K4me3) and the acetylation of lysine 9 of histone H3 (H3K9ac) or lysine 16 of histone H4 (H4K16ac) – are similar genome-wide yet gene orthologs have distinct chromatin and expression patterns. While both *Ogataea* genomes organize into a Rabl conformation, the *O. haglerorum* translocation alters subtelomeric chromatin composition and expression of genes affected by the translocation. Our work highlights the genome function differences that occur on a microevolutionary scale.

**Article Summary:** To assess changes in chromatin – the DNA-protein aggregate controlling genome function – between closely related species of fungi, we compared histone modifications, gene expression, and DNA folding in two yeasts in the same species complex, *Ogataea polymorpha* and *Ogataea haglerorum*. Globally, we found similar patterns of chromatin composition, transcription, and genome folding, with distinct differences in individual genes. Further, a large genome rearrangement in *O. haglerorum* alters histone mark deposition and gene expression at affected chromosome ends. Our results provide insights into functional genome changes that occur over short evolutionary time scales and in response to large chromosomal changes.

## Introduction

In eukaryotes, nuclear DNA associates with proteins to form chromatin, and the composition of this chromatin provides essential control of DNA-templated nuclear processes, including transcriptional gene regulation (Allis and Jenuwein 2016; Courtney et al. 2020). The fundamental unit of chromatin is the nucleosome, in which ~146 basepairs (bp) of DNA is wrapped around an octamer of histone proteins – two each of H2A, H2B, H3, and H4 (Luger et al. 1997; Campos and Reinberg 2009; Luger et al. 2012). The disordered N-terminal histone tails emanating from the core nucleosome particle contain post-translational modifications (PTMs) that provide epigenetic information about the associated DNA (Strahl and Allis 2000; Jenuwein and Allis 2001). Specifically, the nucleosomes in the transcriptionally active euchromatic regions of the genome are marked by the tri-methylation of lysine 4 on histone H3 (H3K4me3), which is enriched at the 5’ start of genes; in addition, multiple lysine residues on each histone protein are hyperacetylated (Strahl et al. 1999; Cheung et al. 2000; Roth et al. 2001; Noma and Grewal 2002; Kurdistani and Grunstein 2003; Ng et al. 2003; Sinha et al. 2006; Freitag 2017). In contrast, the nucleosomes in silent, heterochromatic genomic loci can be methylated on lysine 9 on histone H3 (H3K9me) but are hypoacetylated due to the action of histone deacetylases (HDACs) and have diminished H3K4me3 (Kurdistani and Grunstein 2003; Tamaru et al. 2003; Grewal and Jia 2007; Shahbazian and Grunstein 2007; Grunstein and Gasser 2013; Freitag 2017).

The chromatin state defined by these histone PTMs three-dimensionally (3D) organizes the DNA in eukaryotic genomes. In most eukaryotes, euchromatin is centrally located in the nucleus yet segregated from heterochromatic loci, such as the centromeres and telomeres of chromosomes, which can associate with the nuclear periphery to effectively compartmentalize active and silent genomic loci (Lieberman-Aiden et al. 2009; Rao et al. 2014; Galazka et al. 2016; Falk et al. 2019; Torres, Reckard, et al. 2023). One prominent attribute of fungal genome organization is the Rabl chromosome conformation, characterized by a single centromeric focus that is independent of several telomere clusters, each of which localize to the nuclear periphery (Galazka et al. 2016; Rodriguez et al. 2022; Torres, Reckard, et al. 2023). The Rabl conformation organizes the genomes of budding yeast *Saccharomyces cerevisiae*, the fission yeast *Schizosaccharomyces pombe*, and filamentous fungi including *Neurospora* crassa and Verticillium dahlia (Duan et al. 2010; Mizuguchi et al. 2014; Galazka et al. 2016; Torres, Kramer, et al. 2023). Indeed, the Rabl conformation has been shown by fluorescence microscopy and/or chromosome conformation capture coupled with high-throughput sequencing (Hi-C) experiments in several fungi (Duan et al. 2010; Mizuguchi et al. 2014; Galazka et al. 2016; Klocko et al. 2016; Tanizawa et al. 2017; Torres, Reckard, et al. 2023).

Hi-C is typified by the ligation of adjacent genomic loci into single DNA molecules which, upon paired-end sequencing, report the contact probability of any two loci across the entire genome (Lieberman-Aiden et al. 2009; Lafontaine et al. 2021). In fungal Hi-C, interchromosomal centromeric interactions, as well as intra- and interchromosomal subtelomeric contacts are among the most prominent genomic interactions (Duan et al. 2010; Mizuguchi et al. 2014; Galazka et al. 2016; Winter et al. 2018; Seidl et al. 2020; Rodriguez et al. 2022; Torres, Kramer, et al. 2023; Torres, Reckard, et al. 2023). However, changes to the chromatin composition in fungi can impact genome organization. For example, telomere clustering in *S. cerevisiae* is decreased when the protein Sir3, which strongly binds deacetylated histone tails, is depleted, while Sir3 overexpression promotes telomere bundling (Ruault et al. 2021). Further, *Neurospora* crassa strains deleted of histone deacetylase genes present with heterochromatic hyperacetylation and have increased contact probability between interspersed heterochromatic loci and centromeres, presumably due to the increased chromatin accessibility afforded by hyperacetylation (Scadden et al. 2023; Ebot-Ojong et al. 2025). As also shown in *N. crassa*, loss of H3K9me3, or its cognate binding partner HP1, decompacts constitutive heterochromatic region boundaries, including at the centromeres, but does not abolish inter-heterochromatic interactions (Galazka et al. 2016). Together, these tantalizing clues suggest chromatin composition in general – and more specifically, global histone acetylation patterns – influences fungal genome organization but few studies have examined this relationship in detail.

Although the Rabl chromosome conformation is conserved across diverse fungal species, the histone PTMs associated with heterochromatin silencing have undergone a substantial transition in budding yeasts. Notably, the genes encoding the methyltransferase Clr4 and the chromodomain protein Swi6, needed for H3K9me2-dependent gene silencing, were lost early in Saccharomycotina (Hickman et al. 2011). In *S. cerevisiae*, heterochromatin is established by Sir proteins, including the NAD^+^-dependent HDAC Sir2 (Hickman et al. 2011), which progressively hypoacetylate histones H3 and H4 to form heterochromatin at telomeres and the *HMR* and *HML* silent *MAT* locus cassettes (Rusche et al. 2003; Gartenberg and Smith 2016). However, this mechanism arose following a whole-genome duplication in *Saccharomyces*, which led to subfunctionalization of the *S. cerevisiae* Sir2 protein from its HDAC paralog Hst1, and Sir3 from its paralog Orc1 (Hickman and Rusche 2007; Hickman et al. 2011). Despite these advances, the location of euchromatic and heterochromatic histone PTMs, as well as the mechanism(s) for heterochromatin formation, have not been extensively characterized in other budding yeasts.

While the evolution of genome organization has been examined at broad evolutionary timescales, with numerous Ascomycete and several Basidiomycete fungal genomes shown to form interchromosomal interactions consistent with the Rabl chromosome conformation (Hoencamp et al. 2021; Torres, Reckard, et al. 2023), the vast majority of genome organization (Hi-C) studies in budding yeasts have been performed in *S. cerevisiae* (Duan et al. 2010; Belton et al. 2015; Lee et al. 2016; Eser et al. 2017; Kim et al. 2017; Lazar-Stefanita et al. 2017; Ruault et al. 2021), highlighting the paucity of (epi)genomic studies in other yeasts. Further, few studies have compared genome organization or chromatin composition between closely related species, especially in non-conventional fungi. While the genomes of closely related organisms may be broadly organized with conserved features, including similar chromosome conformations, subtle changes in the hierarchical chromosome organization or distinct changes in local chromatin composition may be apparent.

*Ogataea polymorpha* is a methylotrophic, homothallic yeast species within the order Pichiales (Opulente et al. 2024). *O. polymorpha* is often used in industrial recombinant protein production due to its robust suite of inducible promoters involved in methanol metabolism (Gellissen and Melber 1996) and is an important model for nitrate assimilation (Gellissen 2002), methanol utilization (Hartner and Glieder 2006), peroxisome biology (Gellissen 2002), and the evolution of mating-type switching (Hanson et al. 2014; Maekawa and Kaneko 2014). *O. polymorpha* is one of four members of a species complex that also includes *O. haglerorum* (Naumov et al. 2017). In general, the ~8.9 Mb *O. haglerorum* and *O. polymorpha* genomes are highly syntenic with ~86% sequence identity, yet large genome rearrangements have been reported in *O. haglerorum* isolates (Hanson et al. 2021). In *O. haglerorum* strain 81-453.3, the left arm of chromosome 1 is inverted and translocated adjacent to the chromosome 6 centromere relative to the *O. polymorpha* karyotype (Hanson et al. 2021). Here we present the comparative epigenomics and genome organization for these two members of the *Ogataea polymorpha* species complex. We report the enrichment of H3K4me3, the acetylation of lysine 9 on histone H3 (H3K9ac), and the acetylation of lysine 16 on histone H4 (H4K16ac) in *O. polymorpha* and *O. haglerorum*. We also assessed the genome-wide transcription patterns in both species with RNA-seq datasets to correlate gene expression with chromatin composition, finding distinct epigenetic and transcriptional patterns at conserved genes and chromosome features. Hi-C datasets from *O. polymorpha* and *O. haglerorum* confirmed that both yeast genomes are organized into the conserved Rabl chromosome conformation; Hi-C also validated the translocation reported in *O. haglerorum*. This translocation impacts the composition of the subtelomeric chromatin within the affected donor and acceptor chromosomes. Our work helps to elucidate the unique features of yeast genomes that can occur on a short evolutionary timescale.

## Materials and Methods

Supplemental Materials and Methods, including detailed protocols, are available in File S1.

### Strains and growth conditions

*Ogataea polymorpha* strain SH4330 (NCYC495 *MATα ura3*) (Satoshi Harashima, Osaka University, Osaka, Japan) and *Ogataea haglerorum* isolate 81-453.3 (Phaff collection, University of California-Davis, CA, USA) were grown from freezer stocks on YPD plates (1% w/v yeast extract, 2% w/v peptone, 2% w/v glucose, 2% w/v agar) at 30 °C and individual colonies were grown in YPD broth on shaking incubator at 30°C.

### Nanopore sequencing and generation of the *Ogataea haglerorum* reference genome

The new *O. haglerorum* reference genome was assembled using long-read Oxford Nanopore Technology sequencing (Nanopore-seq), corrected using the previously deposited *O. haglerorum* reference genome (National Center for Biotechnology Information [NCBI] Genome Assembly number ASM1920728v1), which was assembled using Illumina short read whole genome sequencing datasets (NCBI Sequence Read Archive [SRA] Accession number SRX10322527)(Hanson et al. 2021), and refined using Hi-C data (below). For Nanopore-seq, the *O. haglerorum* 81-453.3 strain was grown overnight in YPD media and genomic DNA was extracted using Qiagen Genomic Tip 100/G Kit according to manufacturer’s instructions. Genomic DNA quantity was evaluated using Qubit 3.0 and Broad Range DNA kit according to manufacturer’s instructions. High quality genomic DNA was prepared for sequencing using Rapid Sequencing gDNA Barcoding kit (Oxford Nanopore Technologies) and sequenced on MinION MK1b 9.4.1 flow cell. Basecalling was performed using Guppy v4.2.2, and sequences were trimmed using PoreChop (https://github.com/rrwick/Porechop) and filtered using NanoFilt (Coster et al. 2018) for reads longer than 1000 base pairs (bp). Genome assembly was performed using SMARTdenovo v3 (Liu et al. 2021). Short-read genome sequencing reads (NCBI SRA accession number SRX10322527) were used for two rounds of long-read genome polishing as follows: short-read genome sequences were aligned to the long-read assembly using BWA mem v0.7.17 (Li and Durbin 2009) with parameters “-M -Y -t 4 -R “@RG\tID:RGID\tSM:Ohag\tPL:illumina\tLB:lib1\tPU:unit1”. Samtools (Li et al. 2009) was used to sort and index the aligned reads, and the alignment was used to polish the long-read assembly twice using Pilon v1.24 (Walker et al. 2014); with only two Pilon correction iterations, this *O. haglerorum* genome assembly may still contain assembly errors. Long-read assembly contigs were ordered and assembled into chromosomes via synteny with the *O. polymorpha* reference genome (NCBI genome accession number GCF_001664045.1) using dGenies (Cabanettes and Klopp 2018); dGenies was also used to create synteny dot plots. Finally, Hi-C data (below) was used to correct minor assembly mistakes (e.g., placing some centromeric sequences assembled in the *O. haglerorum* short read datasets but not in long-read datasets) and to properly position the reported translocation between *O. haglerorum* chromosomes 1 and 6, which produced the reference genome reported here. The revised *O. haglerorum* reference genome has been deposited to the NCBI Genome database under Accession Number JBZEMS000000000, and we have provided the *O. haglerorum* reference genome fasta and gff files in Files S2 and S3, respectively. Statistics for the *O. polymorpha* and *O. haglerorum* genomes were generated by QUAST (Gurevich et al. 2013). Genome annotations were predicted by Augustus v3.5.0 (Stanke et al. 2004) with parameters “--species=lodderomyces_elongisporus --strand=both” and tRNAscan-se v2.0.12 (Chan and Lowe 2019). The General Feature Format (GFF) file for displaying *O. haglerorum* genes was created based on the *O. polymorpha* GFF file using LiftOff (Shumate and Salzberg 2021). The *O. polymorpha* Gene Transfer Format (GTF) file was created by converting the *O. polymorpha* GFF file to GTF format using the AGAT software suite (Dainat et al. 2026). Genome synteny was assessed by mapping the *O. polymorpha* and *O. haglerorum* fasta files with minimap2 (Li 2018), output sam files were used in SyRi (Goel et al. 2019) to identify syntenic regions and rearrangements, and chromosomal synteny was plotted using PlotSR (Goel and Schneeberger 2022).

### Chromatin Immunoprecipitation-sequencing (ChIP-seq) library preparation and bioinformatic analyses

ChIP-seq libraries were prepared as previously described (Scadden et al. 2023), except that formaldehyde crosslinked *O. polymorpha* and *O. haglerorum* log-phase cell pellets were used; log-phase cells were obtained from overnight cultures backdiluted into YPD media and grown for ~five to six hours at 30°C until an OD_600_ of 0.8-1.2 was reached. Briefly, resuspended cell pellets were lysed with concurrent chromatin shearing by a Bioruptor Pico, and cleared cell lysate was incubated with one of the following antibodies overnight at 4°C: α-H3K4me3 (ActiveMotif cat# 39060, lot# 31420006); α-H3K9ac (ActiveMotif cat# 39137, lot# 28720002), and α-H4K16ac (ActiveMotif cat# 39068, lot# 08719003). Antibody/histone PTM/DNA complexes were bound by Protein A/G magnetic beads (ThermoFisher cat# PI78609), extensively washed, and the associated DNA was decrosslinked, treated with proteinase K, organically extracted, ethanol precipitated, and Qubit 3.0 quantified. ChIP-DNA sequencing libraries were generated with the New England Biolabs NEB Next Ultra II Illumina barcoding kit, per the manufacturer’s protocols, except that eight PCR cycles were used for library amplification to minimize adenine/thymine (AT)-rich sequence depletion (Ji et al. 2014). ChIP-seq libraries were sequenced at the University of Oregon (UO) Genomics and Cell Characterization Core Facility (GC3F) on an Illumina NovaSeq 6000 as single 118 nucleotide reads (SR118 reads). For display of ChIP-seq enrichment on IGV, fastq files were mapped to the *O. polymorpha* reference genome (NCBI genome accession number GCF_001664045.1), or the newly assembled *O. haglerorum* reference genome (above) using bowtie2 (Langmead and Salzberg 2012). Output sam files were converted to bam files, sorted, and indexed using samtools (Li et al. 2009). RPKM-normalized bigwig files, and subsequent average enrichment profiles over genes, were created using deeptools (Ramírez et al. 2016) for display on Integrative Genomics Viewer (IGV) (Robinson et al. 2011); bed files with all genes, genes in low/high quintiles, or genes in syntenic regions were created and modified in Microsoft Excel. For the box and whisker plots showing the distribution of per-gene average ChIP-seq enrichment, the matrix with bin enrichment values was exported using deeptools and Excel was used to calculate the average enrichment signal for each gene and generate box and whisker plots. JASP (JASP 2026) was used to perform Mann-Whitney tests to compare differences between each independent group of ChIP-seq enrichment distributions.

### Poly-adenine messenger RNA-sequencing (RNA-seq) library preparation and bioinformatic analyses

Total RNA was extracted from *O. haglerorum* log-phase cells, collected similarly to ChIP-seq cell pellets, using a hot acid phenol:chloroform extraction protocol. Briefly, log-phase cell pellets were vortexed with acid phenol, total RNA was ethanol precipitated and DNaseI treated, purified with the RNA Clean and Concentrator-25 kit (Zymo Research), and RNA quantity and quality was evaluated by a Qubit 3.0 (ThermoFisher) and TapeStation (Agilent), respectively. PolyA RNA-sequencing libraries were prepared by the Genomics and Microarray Core at the University of Colorado Anschutz Medical Campus using the Universal Plus mRNA-Seq library preparation kit (Tecan), and 150 bp paired-end sequencing was performed using an Illumina NovaSeq 6000. Published *O. polymorpha* RNA-seq data (NCBI SRA Bioproject Accession# PRJNA418110) (Hanson et al. 2017) were reanalyzed for *O. polymorpha* gene expression. To display species-specific gene expression, the three RNA-seq fastq replicate files were merged together and mapped to the *O. polymorpha* or *O. haglerorum* reference genomes with hisat2 v2.2.1 (Kim et al. 2019); output sam files were converted to bam files, sorted, and indexed with samtools (Li et al. 2009), and Bins Per Million (BPM; the equivalent of Transcripts per Million [TPM])-normalized bigwig files were made by deeptools (Ramírez et al. 2016) for display on IGV (Robinson et al. 2011). For making quintile or top/bottom 5% expressed gene files, each RNA-seq replicate was separately mapped to its respective genome using hisat2 and gene count files for each replicate were generated by htseq (Putri et al. 2022) using the “count” subprogram. The resulting gene count files were normalized for gene length and library size by TPM counts, and the TPM values for each replicate were averaged, in Excel. Genes falling in the highest or lowest quintile (or within the top/bottom 5%) of TPM counts were retained and resulting txt files were made into bed files by changing the file extension. The Pearson correlation of the TPM values for each gene within RNA-seq replicates were calculated in Excel, which showed that all RNA-seq replicates were highly correlated (*O. polymorpha* rep1 vs. rep2 r=0.998874, rep2 vs. rep3 r= 0.991332, and rep1 vs. rep3 r= 0.994489; *O. haglerorum* rep1 vs. rep2 r=0.996014, rep2 vs. rep3 r= 0.988638, rep1 vs. rep3 r=0.981915). To evaluate the significance of the median expression of centromeric or telomeric genes within each species, relative to the expression of all genes within that species, a Mood median test followed by post hoc pairwise Mood tests were performed to obtain FDR-adjusted p-values that indicated significantly different values of medians between bins.

### Chromatin Conformation Capture with high-throughput sequencing (Hi-C) library preparation and bioinformatic analyses

Hi-C libraries were prepared as previously described (Rodriguez et al. 2022) but modified for yeasts using a protocol for *S. pombe* (Tanizawa et al. 2017). Briefly, the nuclei from crosslinked log-phase cell pellets containing ~3.5µg DNA were isolated and made porous, the chromatin was digested by *DpnII* (NEB cat# R0543L), end-repaired with Klenow (NEB cat# M0210L) and dNTPs, including biotin-14-dATP (Invitrogen cat# 19524-016), and ligated with T4 DNA Ligase (NEB cat# M0202L). The ligated DNA was purified, treated with T4 DNA polymerase (NEB cat# M0203L) to remove biotinylated nucleotides from unligated strands, sheared with a QSonica sonicator (model# Q55), and DNA ligation junctions were purified with Streptavidin M280 Dynabeads (ThermoFisher cat# 11205D). Hi-C libraries were barcoded for Illumina sequencing using the NEB Next Ultra II kit for Illumina per the manufacturer’s protocols, except that libraries were amplified with eight PCR cycles to minimize AT-rich sequence depletion (Ji et al. 2014). All Hi-C libraries were sequenced at the UO GC3F on either an Illumina HiSeq 4000 as paired end 100 (PE100) reads or an Illumina NovaSeq 6000 as paired-end 59 nucleotide (PE59) reads.

To display contact probability between genomic loci, high-throughput sequencing fastq files of each species were merged, and the *O. polymorpha* merged Hi-C dataset was mapped to the *O. polymorpha* reference genome (NCBI genome accession number GCF_001664045.1) or the *O. haglerorum* merged Hi-C dataset was mapped to the *O. haglerorum* (above) reference genome using bowtie2 (Langmead and Salzberg 2012) using the --local and --reorder flags. Output sam files were used to build Hi-C contact probability matrices over 1 kilobase (kb) bins with the hicBuildMatrix command of hicExplorer (Ramírez et al. 2018); this program suite which was used for all Hi-C analysis. The command hicMergeMatrixBins generated lower resolution (5 kb or 10 kb) contact probability matrices, while hicPlotMatrix visualized the contact probability over the *O. polymorpha* or *O. haglerorum* genomes, chromosomes, or specific loci. The command hicCorrelate plotted scatterplots and calculated the Pearson correlation values of Hi-C replicates, while the commands hicFindTADs (with flags --correctForMultipleTesting fdr [false discovery rate], --thresholdComparisons 0.3, and --delta 0.15) and hicPlotTADs visualized TAD-like structures across *Ogataea* chromosomes. To directly compare a chromosome 4 syntenic region from *O. polymorpha* and *O. haglerorum*, the command hicNormalize (“multiplicative” option) was used to downscale the values in the *O. polymorpha* contact probability matrix to match the number of *O. haglerorum* valid reads, the command hicAdjustMatrix was used to select the first Megabase of the fourth chromosome from both *Ogataea* matrices, and the commands hicCompareMatrices and hicPlotMatrix were used to display the log_2_ fold change in Hi-C contact probability.

## Results

### Chromatin enrichment and gene expression profiles in two closely related *Ogataea* species

Chromatin structure and genome organization in the budding yeast *Saccharomyces cerevisiae* and the fission yeast *Schizosaccharomyces pombe* are well-characterized, however genome function across a diverse range of fungal species requires further exploration. Here, we compared chromatin composition and genome organization in two closely related yeasts, *O. polymorpha* and *O. haglerorum*, which diverged from *S. cerevisiae* more than 200 million years ago (Shen et al. 2018). The *O. polymorpha* and *O. haglerorum* species were established after the loss of H3K9me2-dependent heterochromatin early in Saccharomycotina evolution but before the expansion of NAD^+^-dependent HDAC genes for regulating heterochromatin, which occurred after the whole-genome duplication event in the *Saccharomyces* lineage (Table S1)(Mannermaa and Oikarinen 1989; Hickman et al. 2011; Grunstein and Gasser 2013; Hanson and Wolfe 2017). This makes species of the *Ogataea* clade attractive candidates to explore novel mechanisms underlying genome function, particularly regarding heterochromatin silencing.

Although the *O. polymorpha* reference genome is nearly complete across its seven chromosomes (Riley et al. 2016), the publicly available *O. haglerorum* reference genome (Phaff strain 81-453.3), assembled from short-read Illumina sequencing, was highly fragmented (NCBI genome assembly ASM1920728v1 statistics: genome size = 8.9 Mb; scaffold # = 64; scaffold N50 = 556.7 kb). To improve the *O. haglerorum* genome and facilitate our analysis of chromatin composition, we performed long-read Oxford Nanopore Technology sequencing of *O. haglerorum* strain 81-453.3, with manual refinement using Hi-C data (below). Our improved *O. haglerorum* genome has seven chromosomes, consistent with *O. polymorpha* (Table S2; genome size = 8.739 Mb; scaffold # = 7; scaffold N50 = 1.286 kb). Comparison of the *O. polymorpha* genome with our new *O. haglerorum* assembly showed that the chromosomes of the two *Ogataea* species are highly syntenic, except for divergent DNA sequences at the chromosome ends (Figure 1A), consistent with previous reports showing that yeast subtelomeres are sites of sequence diversity (Yue et al. 2017). Moreover, comparisons of the *Ogataea* genome sequences confirmed the previously reported translocation between *O. haglerorum* chromosomes 1 and 6 (Figures 1B, S1)(Hanson et al. 2021). Together, this translocation and the reported sequence divergence between these two *Ogataea* species (Hanson et al. 2021) provides an excellent system to study how syntenic or DNA sequence changes affect genome function, including chromatin composition. We performed Chromatin Immunoprecipitation-sequencing (ChIP-seq) for the activating histone PTMs H3K4me3, H3K9ac, and H4K16ac to characterize the genome-wide PTM distribution in *O. polymorpha* and *O. haglerorum*. We chose two acetyl marks and one tri-methylation mark since the loss of H3K9me in budding yeasts, including the *Ogataea* clade, suggests histone deacetylation may be critical for silencing genomic loci. The two replicate ChIP-seq datasets for each histone PTM are highly reproducible (Figure S2), allowing us to merge replicates to display the average histone PTM enrichment. The deposition of each histone PTM is unique from the DNA input into the ChIP reactions for all ChIP-seq datasets (Figure S3), highlighting the specificity of each histone PTM enrichment pattern. To understand the global histone PTM distribution within each *Ogataea* species, we plotted the H3K4me3, H3K9ac, and H4K16ac ChIP-seq enrichment across all genes (Figure 2A). Consistent with its placement in other species, H3K4me3 is primarily enriched at the 5’ side of genes surrounding the transcription start site (TSS) in both *O. polymorpha* and *O. haglerorum* (Figure 2A), although H3K4me3 appears to be enriched across the entire coding sequence of some genes; this extended H3K4me3 enrichment most likely reflects its deposition at small genes, as H3K4me3 peaks are of a uniform size when ChIP-seq enrichment heatmaps are aligned to the TSS as a reference point (Figure S4). In contrast, both acetyl marks in each species are primarily enriched over gene bodies but are reduced at the TSS and transcription termination sites (TTS), although these boundaries are less clear in *O. haglerorum*. However, some genes in each species had minimal histone PTM enrichment (Figure 2A, plot bottoms). We speculated that this reduced histone PTM deposition may be connected to gene expression. To address if histone PTM enrichment and transcription levels could be correlated, we performed polyadenosine messenger RNA sequencing (mRNA-seq) in *O. haglerorum* and analyzed published *O. polymorpha* mRNA-seq data (Hanson et al. 2017). We divided the gene expression counts into quintiles and plotted the average histone PTM enrichment over the lowest and highest expressed gene quintiles. In *O. polymorpha*, the lowest expressed genes have minimal H3K4me3 enrichment at the TSS while H3K9ac is substantially decreased and H4K16ac is maintained (Figure 2B, left). *O. haglerorum* similarly has minimal H3K4me3 enrichment over the lowest gene quintile, along with decreased H3K9ac, while H4K16ac is mostly preserved (Figure 2B, right). In contrast, the genes in the highest expression quintile in both species have extensive H3K4me3 and histone hyperacetylation (Figure 2B). We conclude that H3K4me3, H3K9ac, and H4K16ac have globally similar enrichment patterns that are correlated to transcriptional levels.

**Figure 1.**
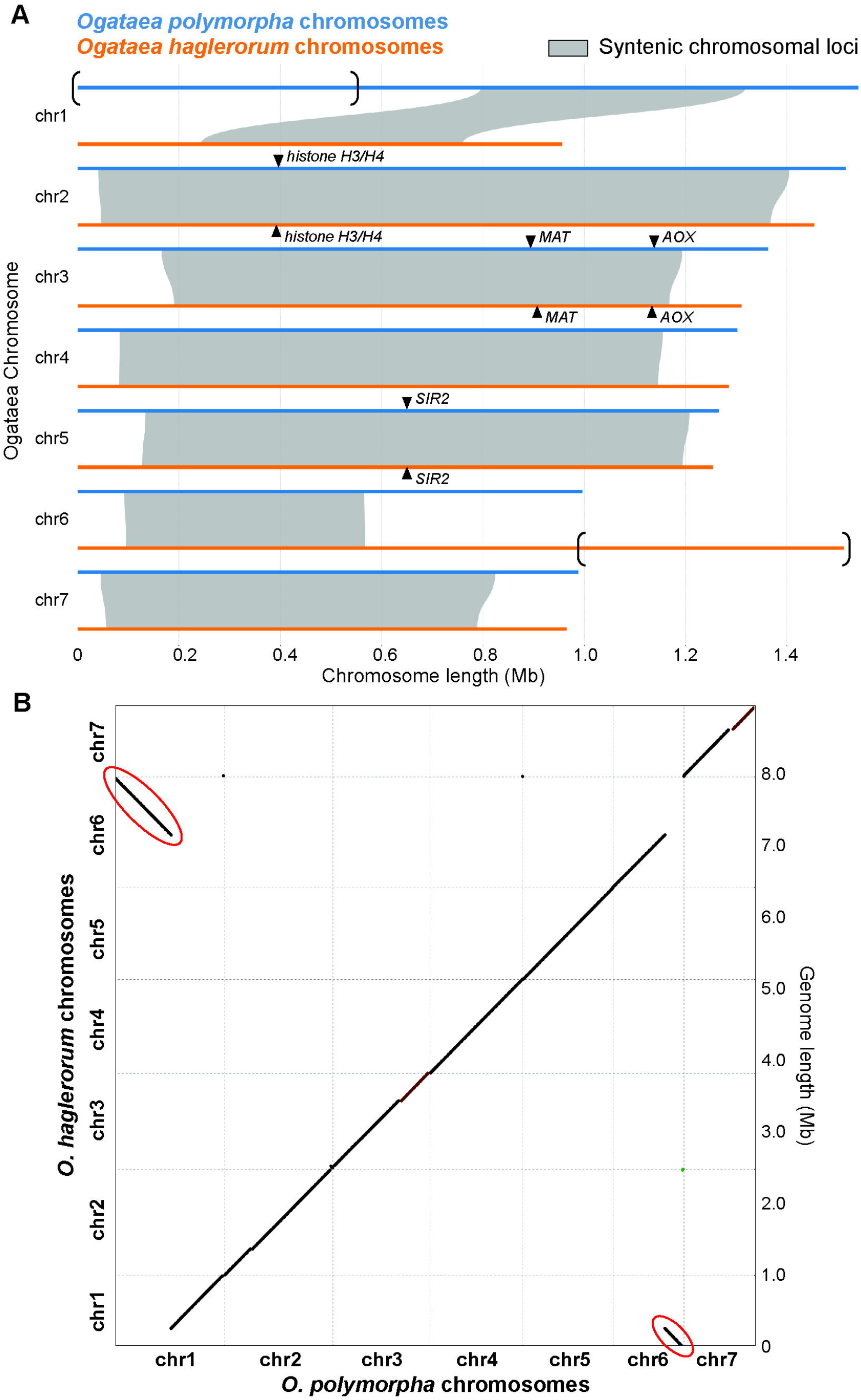
The chromosomes in two *Ogataea* species are highly syntenic. (A) Synteny plots showing the sequence conservation between *O. polymorpha* chromosomes (blue lines) and *O. haglerorum* (orange lines) for all chromosomes. Gray shading shows syntenic loci. Individual genes subject to chromatin inspection are indicated with black arrowheads. (B) dGenies dot plot of the synteny between the *O. polymorpha* (x-axis) and *O. haglerorum* (y-axis) chromosomes. Dark brown lines indicate identity (BLAST-like alignment) values of the syntenic chromosomes between 0 to 0.25, while light brown lines highlight identity values between 0.25 to 0.5 (Cabanettes and Klopp 2018). Red ovals highlight the region of the *O. polymorpha* chromosome 1 that is translocated onto *O. haglerorum* chromosome 6, as previously reported (Hanson et al. 2021) and confirmed by Hi-C (this work).

**Figure 2.**
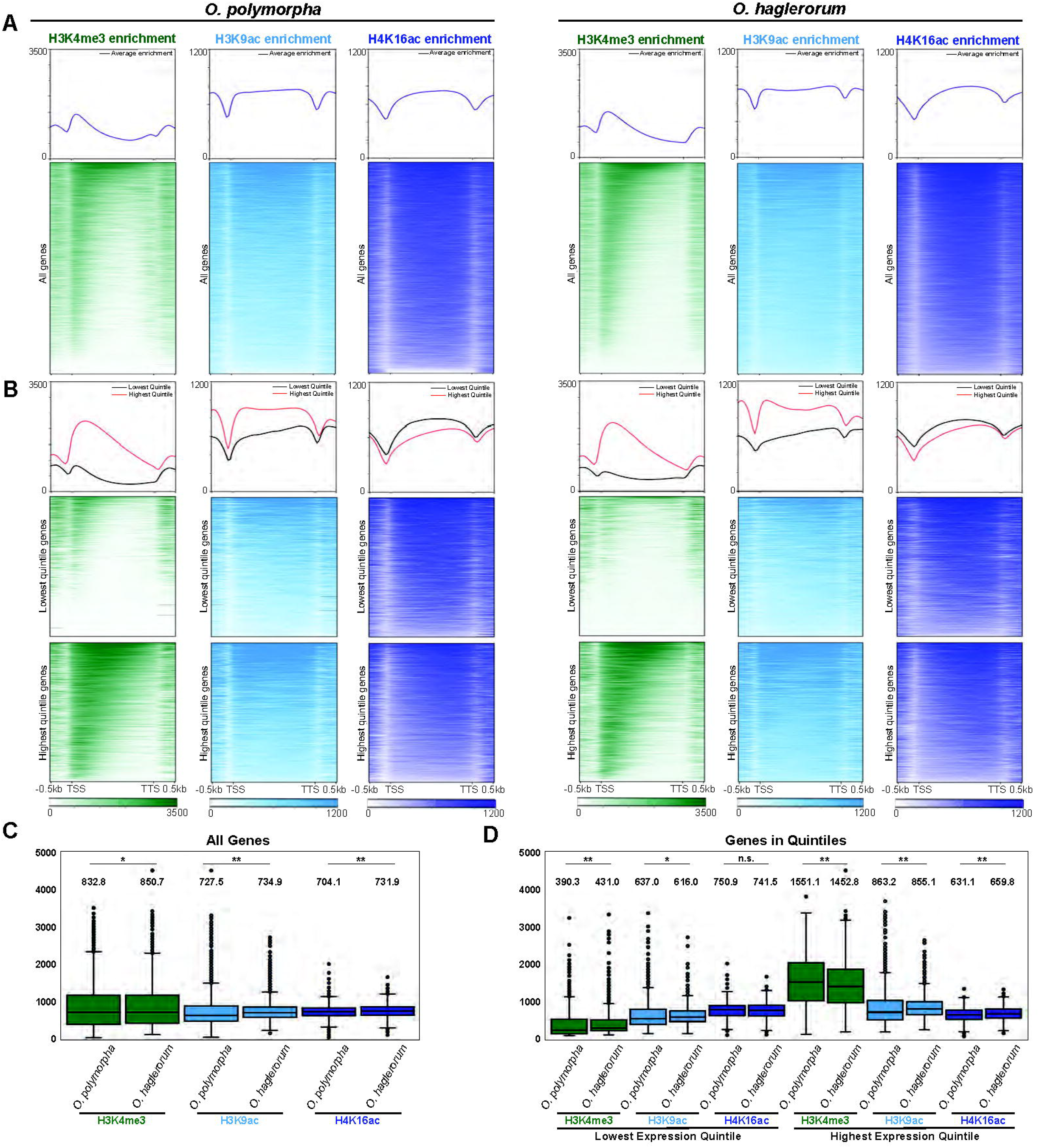
Conserved global gene enrichment patterns for three histone PTMs in two *Ogataea* species. (A-B) Average enrichment profiles (top images) and heatmaps (bottom images) of H3K4me3 (green), H3K9ac (light blue), and H4K16ac (dark blue) over (A) all *Ogataea* genes or (B) the lowest (top) or highest (bottom) quintile of expressed genes. Each gene is scaled to two kilobases, and 500 bp is plotted before and after the Transcription Start Site (TSS) and Transcription Termination Site (TTS). ChIP-seq enrichment value scalebars are shown on the bottom. The left three histone PTM enrichment profiles/heatmaps correspond to genes in the *Ogataea polymorpha* genome, while the right three profiles/heatmaps show the patterns for the *Ogataea haglerorum* genome. (C-D) Box and whisker plots showing the distribution of the average histone PTM ChIP-seq enrichment over (C) all genes or (D) low and high quintiles in both *Ogataea* species. The average ChIP-seq enrichment of the gene average within that group is shown above each plot. Asterisks indicate significantly different distributions of the ChIP-seq enrichment between the *Ogataea* species (Mann-Whitney test; ** = p-value < 0.001, * = p-value < 0.01, n.s. = not significant).

Next, we determined if gene expression and histone PTM distribution are correlated with conserved genome features by characterizing the chromatin profiles around centromeres and in subtelomeric regions, both of which typically show reduced transcription. We first examined the expression of genes near these two chromosomal feature*S. O. polymorpha* and *O. haglerorum* centromeres span ~10 to 20 kb and contain long-terminal repeats (LTRs) and other retroelement derived sequences (Ravin et al. 2013; Hanson et al. 2014; Hanson et al. 2021). Overall, genes within 10 kb of the centromere in both *O. polymorpha* and *O. haglerorum* were not reduced in expression relative to the expression level of all genes (Figure 3A). In contrast, subtelomeric genes within 10 kb of the telomere repeats showed statistically significant lower expression levels (Figure 3A). The average histone PTM enrichment in both species is consistent with these expression patterns, as H3K4me3 deposition occurs over the TSS of genes within 10 kb of the centromeres but H3K4me3 at the TSS of genes within 50 kb of the telomeres is reduced (Figure 3B); here, we designated the chromatin within 50 kb of the chromosomes ends as the “subtelomeres” to be consistent with the clustering of chromosome ends, as elucidated by *Ogataea* Hi-C data (below). Also, histone acetylation is moderately enriched across pericentromeric and subtelomeric gene bodies in both species (Figure 3B). Histone PTM enrichment and transcription levels surrounding individual chromosome features support the broader trends found in the average profiles. To this end, we displayed H3K4me3, H3K9ac, and H4K16ac ChIP-seq enrichment tracks, as well as a track of the average, BPM (TPM)-normalized mRNA-seq reads, from both species. As shown in Figure 3C, while the chromosome 5 centromeres differ in length in *O. polymorpha* and *O. haglerorum* (~18 kb and ~10 kb, respectively), the centromeres of both species show hallmarks of heterochromatin silencing, with regions devoid of H3K4me3 and histone acetylation. Strikingly, a Ty-like retrotransposon in the *O. polymorpha* chromosome 5 centromere appears to be lowly transcribed and is marked with euchromatic histone PTMs (Figure 3C). While all other *O. polymorpha* and *O. haglerorum* centromeres have similar heterochromatic profiles characterized by reduced histone acetylation and H3K4me3, the *O. polymorpha* centromeres on chromosomes 1, 3, and 7 also contain retroelements (Figures S5-S6). In addition, the pericentromeric chromatin in each species is quite variable, with genes in this region showing evidence of transcription and enrichment of histone PTMs (Figures 3C, S5-S6). Conversely, the subtelomeres of both species have hallmarks of silent heterochromatin. As shown for chromosome 6 of *O. polymorpha* and *O. haglerorum*, the left and right subtelomeres harbor numerous genes within the lowest expression quintile that exhibit a histone PTM profile consistent with heterochromatin, including reduced histone acetylation and few TSS peaks of H3K4me3, although H4K16ac deposition is still prevalent (Figure 3D). Similar heterochromatic chromatin patterns occur over the subtelomeres of the other six chromosomes in both species (Figures S7-S8). Notably, the heterochromatic region of the *O. haglerorum* chromosome 6 right subtelomere is smaller than its *O. polymorpha* counterpart, as only ~eight repressed genes occur immediately adjacent to the *O. haglerorum* telomere before the chromatin transitions into highly expressed euchromatin (Figure 3D). Together, we conclude that the centromeres and subtelomeres of both *Ogataea* species are heterochromatic, given the histone hypoacetylation, reduced H3K4me3, and lower gene expression across these chromosome features, but that pericentromeric regions in *Ogataea* species are not heterochromatic, consistent with other yeasts (Kapoor et al. 2015; Sreekumar et al. 2019).

**Figure 3.**
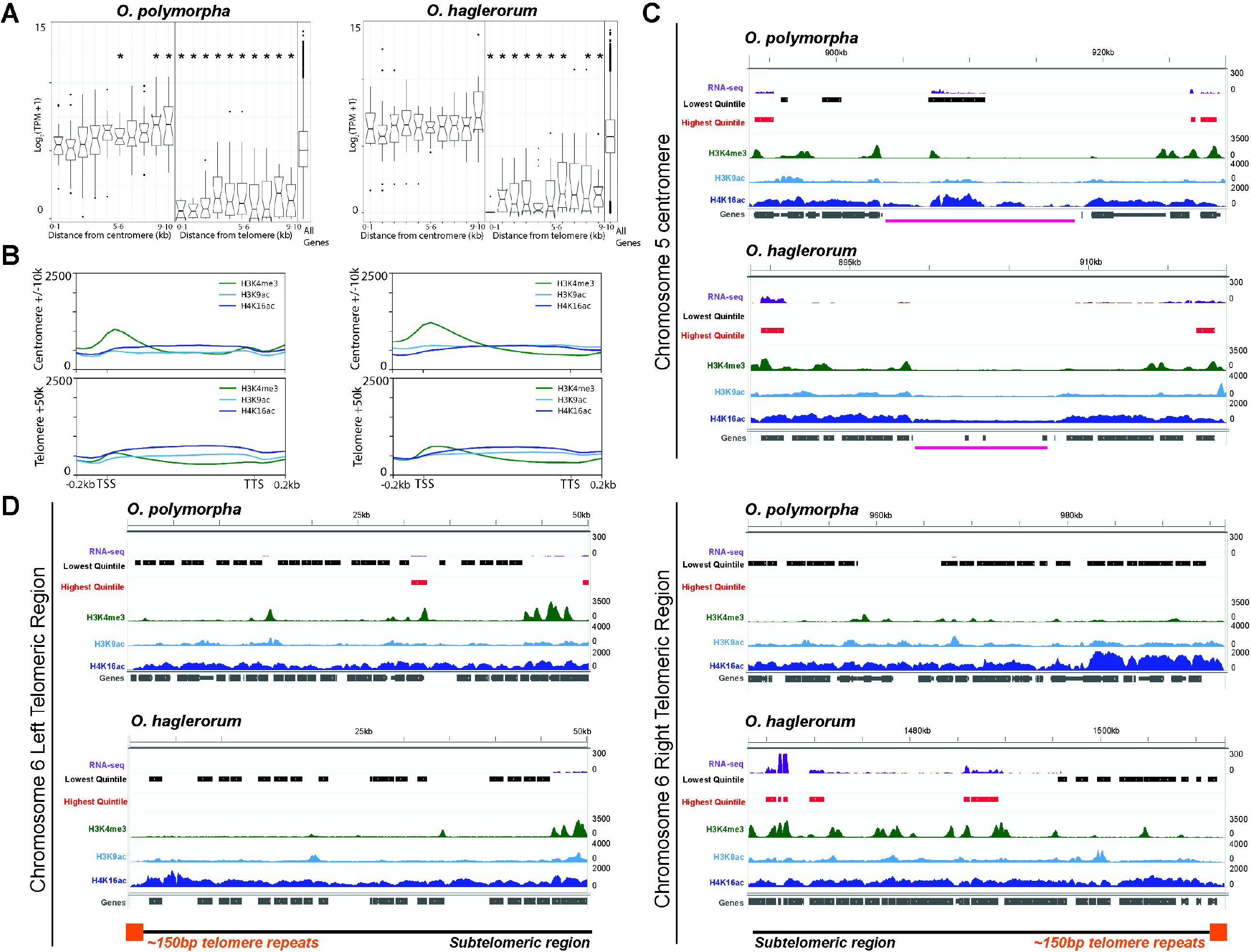
Conserved centromeric and telomeric flanking chromatin composition and gene expression patterns in two *Ogataea* species. (A) Box and whisker plots of the median expression of genes binned into 1 kb segments within the 10 kb flanking all seven centromeres (left side of each plot) or all 14 telomeres (right side of each plot); median expression of all genes is shown on the right end of the plot. Significant differences between the centromeric or telomeric bin median and the All Gene bin median are indicated by asterisks (FDR-adjusted p-value < 0.05; pairwise Mood median tests). (B) Average enrichment profiles of three histone PTMs, from the Transcription Start Site (TSS) to the Transcription Termination Site (TTS) of the genes that are found within 10 kb of all centromeres or genes that are found within 50 kb of all telomeres for *O. polymorpha* (left panels) or *O. haglerorum* (right panels). Each histone PTM is colored and scaled as in Figure 2. (C-D) IGV images of histone PTM enrichment tracks of H3K4me3 (green), H3K9ac (light blue), and H4K16ac (dark blue), RNA-seq gene expression values (purple), or genes (gray boxes) across the (C) chromosome 5 centromere or (D) the chromosome 6 left and right subtelomeres, for *O. polymorpha* (top images) or *O. haglerorum* (bottom images). Centromeres in panel C are indicated by pink lines, while telomere repeats in panel D are indicated by orange boxes. Genes in the lowest expression quintile are marked by black lines while genes in the highest expression quintile are indicated by red lines.

To determine if the chromatin composition patterns are also broadly maintained across individual genes, we examined histone PTM enrichment and transcription across specific, syntenic genes in *O. polymorpha* and *O. haglerorum* (Figure 1A) that are either highly or lowly expressed. As an example of highly transcribed genes, we examined the genes encoding histones H3 and H4 in *O. polymorpha* and *O. haglerorum*, which are in a divergent orientation (Figure 4A), similar to that of other eukaryotes (Smith and Andrésson 1983; Woudt et al. 1983; Bernhard and Schlegel 1998; Hays et al. 2002). The highly expressed *Ogataea* histone genes have distinct patterns of activating histone PTM enrichment in each *Ogataea* species, with *O. haglerorum* having greater enrichment of H3K4me3 relative to *O. polymorpha*, but surprisingly, the histone genes in each species are similarly hypoacetylated (Figure 4A). As an example of a lowly transcribed gene, the methanol catabolism gene *AOX*, which allows growth in methanol-containing media, has minimal H3K4me3 and H3K9ac enrichment (Figure 4B), indicative of a more repressed chromatin profile, which is consistent the rich media growth conditions used for ChIP-seq (see Materials and Methods). We also examined two additional genes that are biologically relevant to *Ogataea* research. First, we examined the chromatin composition at the *MAT* locus, since previous work showed *O. polymorpha* uses “flip-flop” mating-type switching, with the *MAT* genes closest to the centromere being silenced (Hanson et al. 2014; Maekawa and Kaneko 2014); we note that both the *O. polymorpha* and *O. haglerorum* genome assemblies place the *MATα* locus proximal to the centromere. Interestingly, only minor chromatin enrichment differences occur over the *MAT* genes of each species (Figure S9). Specifically, in *O. polymorpha*, the *MATα* genes have an active chromatin composition while the histone PTMs over the *MATa* genes have a silent chromatin composition, consistent with the *MATα* mating-type of the SH4330 strain (Figure S9A, left). However, in *O. haglerorum*, both the *MATa* and *MATα* genes are moderately enriched with histone acetylation and H3K4me3, reflecting a mixed cell population undergoing mating-type switching, as confirmed by PCR (Figures S9A, right; S10). Finally, examination of the *SIR2* gene, which encodes a moderately-expressed histone deacetylase that could be essential for silencing chromatin in *Ogataea* species, showed that similar H3K4me3 peaks occur over the TSS and that gene body histone hyperacetylation is present in both species, although the peaks of H3K9ac are unique (Figure S9B).

**Figure 4.**
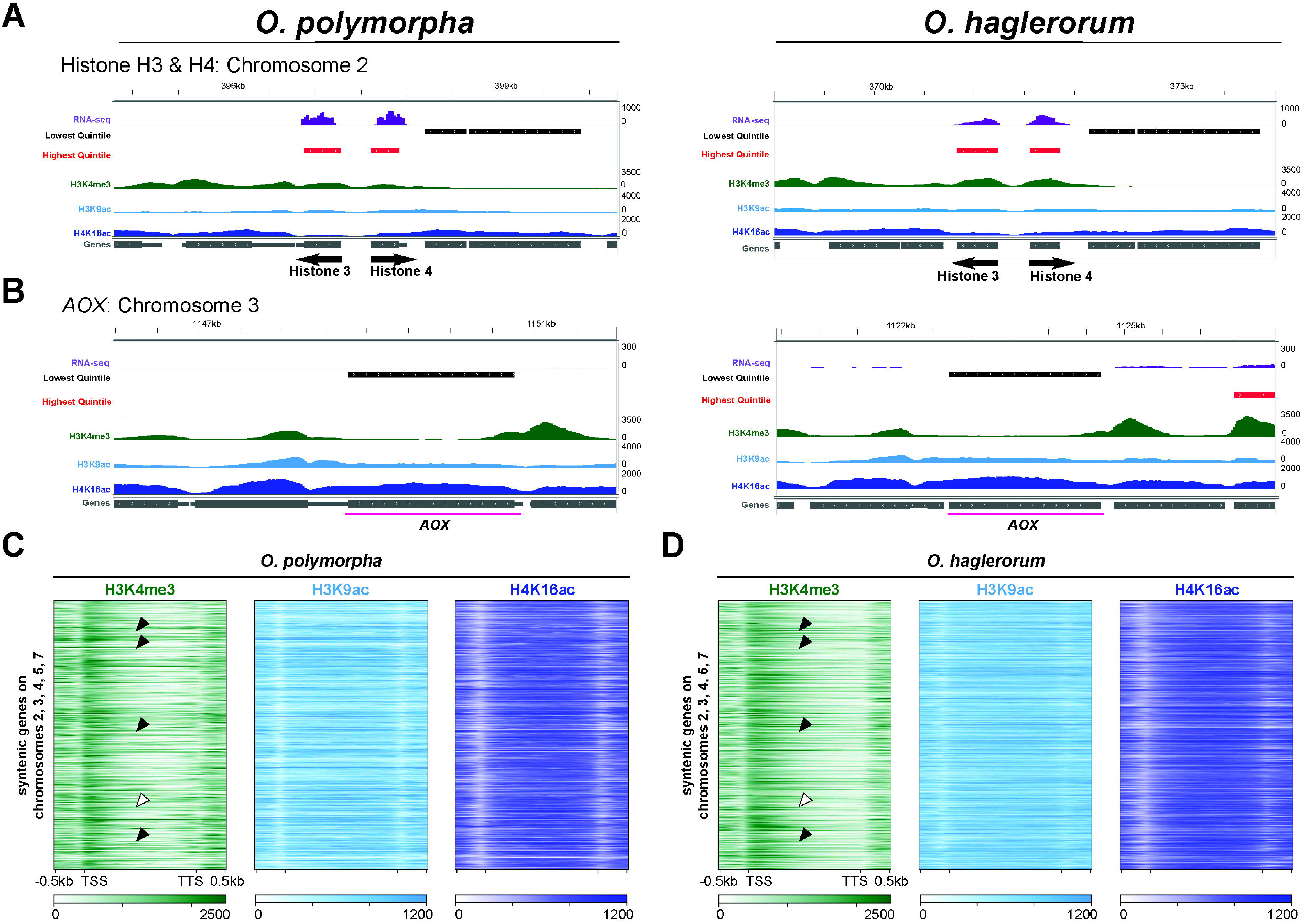
Distinct expression levels and chromatin composition patterns in *O. polymorpha* and *O. haglerorum* genes. (A-B) IGV images of histone PTM enrichment and gene expression values, plotted as in Figures 3C-D, over select genes, including (A) the highly expressed bidirectional histone H3/H4 genes and (B) the lowly expressed *AOX* alcohol oxidase gene. (C-D) Average enrichment heatmaps of histone PTM ChIP-seq enrichment, plotted similarly to those in Figure 2A, except that only syntenic genes on chromosomes 2, 3, 4, 5, and 7 are shown; the gene order is maintained in all plots. Black arrowheads indicate examples of species-specific chromatin patterns, while the white arrowhead shows an example of a gene with a similar chromatin enrichment pattern.

To understand if differences in chromatin composition of individual genes reflect a broader pattern across all of the genes within syntenic regions of the *O. polymorpha* and *O. haglerorum* genomes, we generated ChIP-seq enrichment heatmaps that maintain the gene order of the primarily syntenic chromosomes 2, 3, 4, 5, and 7; genes in the divergent chromosomes 1 and 6, as well as the subtelomeres of all chromosomes were excluded. As shown in Figures 4C-D, the histone PTM patterns in *O. polymorpha* and *O. haglerorum* are qualitatively distinct, suggesting gene-specific chromatin composition differences are widespread in *Ogataea* clade species. In fact, quantitative assessment of the *O. polymorpha* and *O. haglerorum* per-gene ChIP-seq enrichment average across all genes, or the lowest/highest quintile genes, showed the distributions of the average histone PTM enrichment across gene groups are statistically distinct in each species, except for H4K16ac in the lowest gene quintile (Figure 2C-D; Mann-Whitney statistical tests), further supporting that the chromatin composition over the genes in each *Ogataea* species is unique. Together, we conclude that the *O. polymorpha* and *O. haglerorum* genomes have broadly comparable chromatin profiles in aggregate gene analyses, across centromeres and telomeres, and a minority of genes. However, gene-specific differences in chromatin composition can occur between each *Ogataea* species.

### *Ogataea* genomes form similar Rabl chromosome conformations but exhibit notable organization differences

Budding yeast and other fungal genomes are often organized into the Rabl chromosome conformation, which is characterized by a single centromere cluster that is independent of telomere bundles, each of which associate with the nuclear periphery (Torres, Reckard, et al. 2023). To understand the organization of the seven chromosomes of the *O. polymorpha* and *O. haglerorum* genomes, we performed chromosome conformation capture with high-throughput sequencing (Hi-C) on both species. We chose the restriction enzyme *Dpn*II (recognition sequence 5’ ^GATC) for our Hi-C experiments since both *Ogataea* genomes have an ~50% GC-content (~47.9% GC-basepairs for *O. polymorpha* and ~49.7% for *O. haglerorum*)(Hanson et al. 2021) and a relatively even distribution of *Dpn*II sites within gene-rich euchromatin (Figure S11). Our Hi-C replicates are well-correlated and replicate contact probability heatmaps of single chromosomes show similar patterns (Figure S12), allowing us to merge both replicates into one Hi-C dataset for each species reflecting the average genome organization across a cell population. The merged *O. polymorpha* Hi-C dataset contains 28,894,823 valid reads (690.7 contacts per *Dpn*II site) while the *O. haglerorum* Hi-C dataset contains 18,600,424 valid reads (433.6 contacts per *Dpn*II site), suggesting that with a high density of Hi-C contacts per *Dpn*II site, each Hi-C dataset would accurately portray the respective genome organization for each species.

We examined the organization of the seven *O. polymorpha* chromosomes using a Hi-C contact probability heatmap, displayed either with raw or Knight Ruiz (KR) corrected values, with the latter used to remove inherent biases in the underlying DNA sequences (Knight and Ruiz 2013). Figure 5A shows the *O. polymorpha* genome is organized into a Rabl conformation, as dark inter-chromosomal centromeric foci with reduced centromere-chromosome arm contacts are observed, consistent with a centromere bundle; we note that the region depleted in the middle of *O. polymorpha* chromosome 7 is the site of the ribosomal DNA repeats and due to their repetitive nature, Hi-C reads cannot be specifically mapped to this locus. Moreover, intra- and inter-chromosomal subtelomere clusters are readily distinguishable in whole genome and two individual chromosome heatmaps (Figures 5A-B). Indeed, Hi-C heatmaps of chromosomes 1 and 6 show strong intra-chromosomal subtelomeric clustering of the left and right chromosome ends (Figure 5B, black arrows). In contrast, few interactions are observed between the centromeres and the surrounding euchromatin within individual chromosomes (Figure 5B). Inter-chromosomal interactions between centromeres or between subtelomeres are apparent in high-resolution (2.5 kb bin) contact frequency heatmaps between chromosomes 1 and 6 (Figure 5C, left). The subtelomeric cluster interactions extend roughly 50 to 100 kb from the telomere repeats (Figure 5C, middle), while the strongest inter-centromeric contacts center around the centromere yet extend ~100 kb across the pericentromeric regions, consistent with this centromere-flanking euchromatin forming “clothespin” structures as this DNA emanates from the centromeric chromatin bundle (Figure 5C, right). Also prominent on single chromosome heatmaps are TAD-like chromatin loop structures immediately off diagonal, which appear as squares of enhanced interactions (Figure 5B). To computationally assess the presence of TAD-like structures, we predicted TAD boundaries for *O. polymorpha* chromosome 6 (Figure 5D). The computationally predicted TAD-like structures agree with our visual inspection, although one TAD spanned the centromere (Figure 5D, black box), which would not be biologically relevant in the Rabl conformation, and highlights the current limitations of fungal TAD prediction. Still, the TAD prediction indicated multiple ~50 kb TAD-like structures on the euchromatic chromosome arms, which appear to be hierarchically nested within several larger chromatin loops (Figure 5D). To characterize the chromatin properties of these TAD-like structures, we plotted the histone PTM ChIP-seq tracks and the highest or lowest 5% of mRNA-seq expressed genes below the Hi-C contact probability data of chromosome 6. Interestingly, strong TAD-like structure boundaries appear to contain highly expressed genes and prominent peaks of H3K9ac (Figure 5D, blue arrows); several small H3K4me3 peaks also overlap with other TAD-like structure boundaries. These observations suggest highly expressed *O. polymorpha* genes could define chromatin loop boundaries. In contrast, the silent genes in subtelomeric regions form isolated structures comprised of strong contacts, despite the absence of a computationally defined TAD-like structure boundary (Figure 5D, purple lines). Together, we conclude that the *O. polymorpha* genome forms a Rabl conformation characterized by bundles of silent chromosome features and nested euchromatic loops.

**Figure 5.**
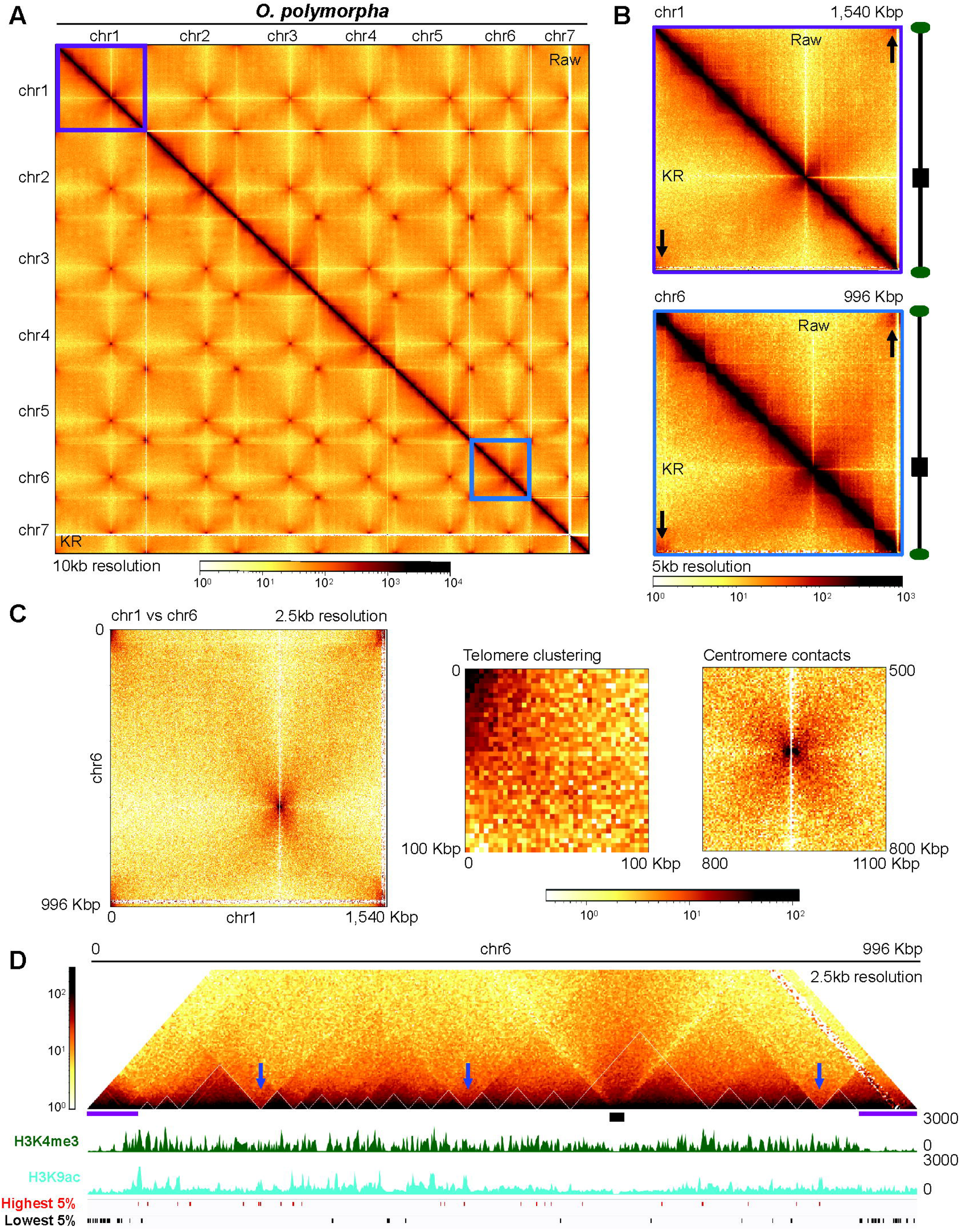
The *O. polymorpha* genome forms a Rabl chromosome conformation characterized by hierarchical chromatin loops. (A-B) Hi-C contact probability heatmaps of (A) the whole *O. polymorpha* genome at 10 kb resolution or (B) chromosome 1 (purple box in A) and chromosome 6 (blue box in A) at 5 kb resolutions. In all Hi-C contact probability heatmaps, raw Hi-C data is shown above the diagonal while the Knight-Ruiz corrected (KR) contact probability heatmap to reduce underlying sequencing biases, is displayed below the diagonal. Scalebars of log_10_ read counts are shown below the heatmaps. In panel B, chromosome schematics with the centromere (black boxes) and telomere (green ovals) positions are shown to the right and arrows indicate the intra-chromosomal subtelomeric contacts. (C) Hi-C contact probability heatmaps of the inter-chromosomal interactions at 2.5 kb resolution between (left) the entire chromosome 1 (x-axis) and chromosome 6 (y-axis); (center) the interactions surrounding the subtelomeres; or (right) the interactions around the centromeres. (D) Hi-C contact probability image of chromosome 6 with TAD-like structures plotted (white triangles). ChIP-seq enrichment tracks for H3K4me3 (green) and H3K9ac (light blue), or the highest (red lines) or lowest (black lines) 5% of genes are plotted below the heatmap. The centromeres (black box) and the subtelomeres (purple lines) on chromosome 6 are indicated. The blue arrows denote strong boundaries of TAD-like chromatin loop structures.

We next examined Hi-C data from *O. haglerorum* to determine if these genome organization properties are conserved between species. When mapping Hi-C reads to the *O. haglerorum* genome with *O. polymorpha* synteny, the previously reported translocation between chromosomes 1 and 6 is readily observed (Figure 6A, top; Figure S13)(Hanson et al. 2021). In *O. haglerorum*, ~585 kb of the left arm of chromosome 1 is flipped and inserted immediately adjacent to the chromosome 6 centromere (Figure 6A, bottom schematic). Guided by these Hi-C data, we rearranged the *O. haglerorum* contigs to account for this translocation (see Materials and Methods) and remapped the *O. haglerorum* Hi-C reads to the corrected reference genome, which restored the contiguous Hi-C diagonal of close proximity contacts (Figure 6B-C). This Hi-C contact probability heatmap also shows the *O. haglerorum* genome is organized into a Rabl conformation, with strong inter-chromosomal centromeric foci independent of subtelomeric bundles (Figure 6B). Individual contact probability heatmaps of *O. haglerorum* chromosomes 1 and 6 show the isolation of centromeres from the gene-rich euchromatin of chromosome arms (Figure 6C), while *O. haglerorum* subtelomeres exhibit intra-chromosomal clustering comparable to *O. polymorpha* (Figure 6C, arrows). High resolution inter-chromosome heatmaps also show the prominent centromere clusters that are independent of subtelomeric bundles (Figure 6D). Closer visual inspection of *O. haglerorum* chromosome 6 shows multiple TAD-like chromatin loop structures within each euchromatic arm, as supported by computational TAD prediction (Figure 6E). These TAD-like structures, including evidence of hierarchical loops, are similar to those observed for the *O. polymorpha* genome. However, the *O. haglerorum* TADs are predicted to be of a smaller size (~20 to 50 kb) relative to those on chromosome 6 in *O. polymorpha*, and strong TAD-like structure borders may be correlated with moderate-sized H3K9ac peaks (Figure 6E, blue arrows), suggesting subtle folding differences exist within the euchromatin of *O. haglerorum* and *O. polymorpha*. Regarding the heterochromatic subtelomeres, the silent *O. haglerorum* subtelomeres also form dense structures at chromosome ends, similar to that of *O. polymorpha* (Figure 6E, purple lines). Together, these Hi-C data suggest that the two *Ogataea* species organize their genomes through broadly conserved mechanisms, although smaller, species-specific chromatin composition and TAD-like structure formation differences exist.

**Figure 6.**
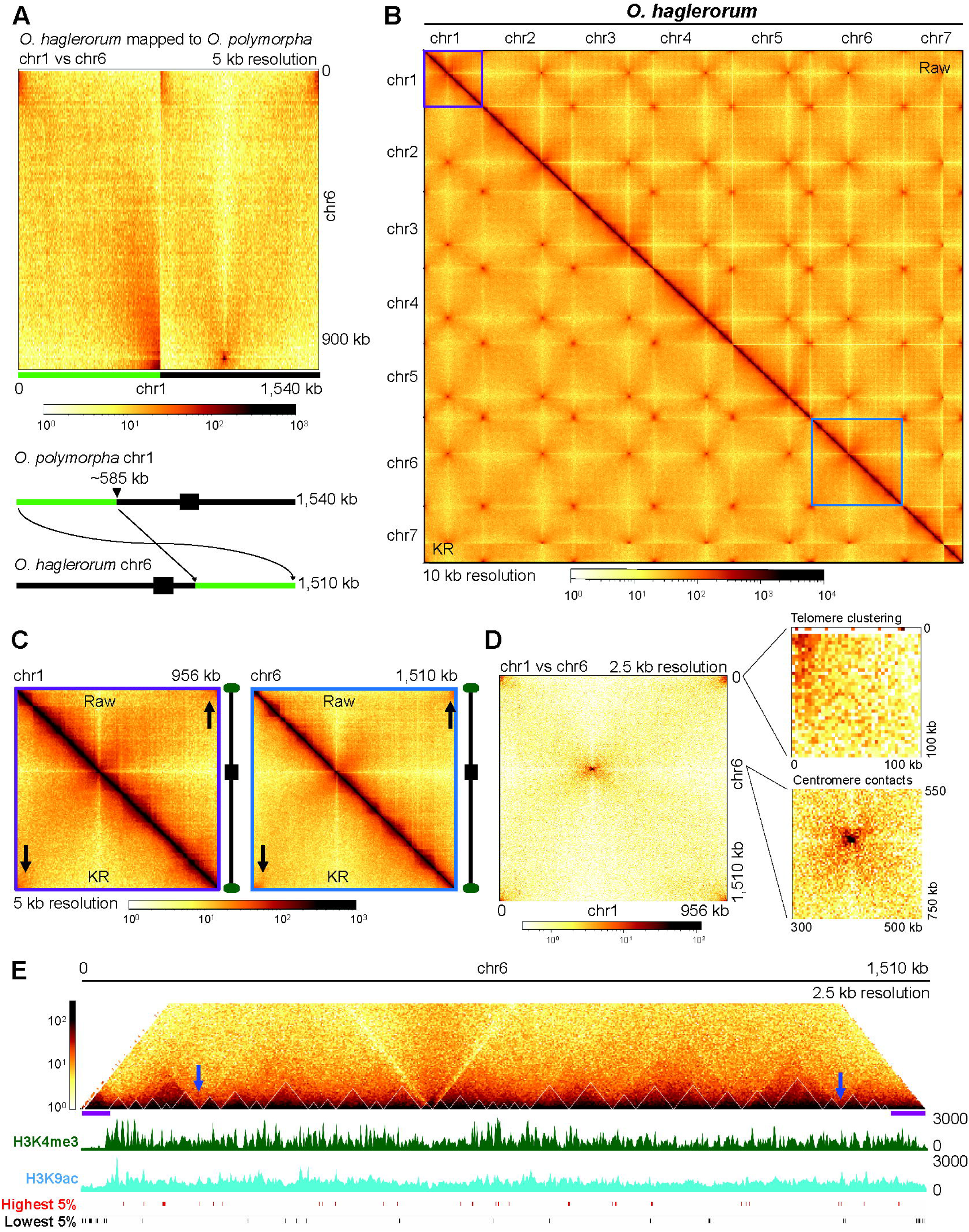
The translocation in the *O. haglerorum* genome alters chromatin loops while maintaining the Rabl conformation. (A) Inter-chromosomal contact probability heatmap of the interactions between chromosomes 1 (x-axis) and 6 (y-axis) at a 5 kb resolution of *O. haglerorum* Hi-C data mapped to a reference genome reflecting the *O. polymorpha* synteny. The scalebar of log_10_ read counts is shown below the heatmap. A schematic of the inter-chromosomal translocation (translocated sequence indicated by the green lines; centromeres by black boxes) that occurred between chromosome 1 (top) and chromosome 6 (bottom) in *O. haglerorum* is shown at the bottom, with matching colored lines shown on the heatmap above. (B-C) Hi-C contact probability heatmaps of (B) the whole *O. haglerorum* genome or (c) chromosomes 1 (purple box in B) or 6 (blue box in B), presented as in Figure 5A-B. (D) Inter-chromosomal Hi-C contact probability heatmaps showing the interactions between the *O. haglerorum* chromosomes 1 (x-axis) and 6 (y-axis)(left), the subtelomeres (top right), or centromeres (bottom right), as in Figure 5C. (E) Hi-C contact probability image of *O. haglerorum* chromosome 6 with TAD-like structures (white lines), plotted as in Figure 5D.

### The *O. haglerorum* genome translocation alters the donor and acceptor chromosome chromatin composition

The translocation between chromosomes 1 and 6 in *O. haglerorum* is the most striking difference between the two *Ogataea* genomes and has the potential to harbor distinct changes to yeast genome function. This translocation occurred uniquely in the *O. haglerorum* strain 81-453.3 and is not shared by any other *O. haglerorum* isolates (Figure S14). Assuming only one DNA break occurred in this rearrangement, the *O. haglerorum* left subtelomere on chromosome 1 had to be reestablished, while the previous right subtelomere on *O. haglerorum* chromosome 6 would localize internally when the chromosome 1 DNA was placed proximal to the chromosome 6 centromere (Figure 6A, schematic). We speculated that the underlying subtelomeric chromatin composition would be altered within the affected chromosomes in both yeast species. To determine if differences in subtelomeric histone PTM enrichment or gene expression are present, we plotted the Hi-C heatmaps of the affected chromosomes, scaled to their respective sizes, and examined the H3K4me3, histone acetylation, and gene expression IGV tracks at the subtelomeres.

Immediately noticeable was the size of the affected chromosomes, as the *O. polymorpha* chromosome 1 is distinctly longer than chromosome 6, while the opposite is true for *O. haglerorum* (Figure 7A-B). Each chromosome has intra-chromosomal subtelomere bundles, suggesting that the *O. haglerorum* chromosome 1 reestablished the left subtelomere post-translocation. Moreover, no chromosome-internal subtelomere contacts are observed in *O. haglerorum* chromosome 6 at a location where the previous subtelomere was positioned prior to the translocation (Figure 7B, bracket), indicating that any remaining chromosome-internal subtelomeric features were resolved. Further, two similar off-diagonal TAD-like structures occur in both *O. polymorpha* chromosome 1 and *O. haglerorum* chromosome 6 (Figure 7A-B, purple lines), although the *O. haglerorum* structures are weaker, implying these TAD-like structures are modular. As a control, we assessed if the first Megabase of the primarily syntenic chromosome 4 of each *Ogataea* species also contains TAD-like structures, and indeed, multiple TAD-like structures are conserved on each chromosome 4 in *O. polymorpha* and *O. haglerorum* (Figure S15). Examination of the subtelomeric chromatin composition shows that each chromosome end has multiple silent genes characterized by reduced H3K4me3 and H3K9ac enrichment and moderate H4K16ac levels, which are maintained despite being involved in the translocation (Figure 7B). For example, the syntenic *O. polymorpha* chromosome 1 left subtelomere and the *O. haglerorum* chromosome 6 right subtelomere have qualitatively similar properties, as the heterochromatin in both species extends only ~10-20 kb beyond the telomere repeats, suggesting the translocation simply moved the subtelomeric chromatin between chromosomes (Figure 7B, top and bottom IGV images). The adjacent euchromatin, with increased H3K4me3 and histone hyperacetylation is also comparable, with multiple highly expressed genes within our arbitrary 50 kb subtelomeric boundaries, although the absolute patterns are not identical (Figure 7B, top and bottom images). Further, the newly formed *O. haglerorum* chromosome 1 left subtelomere strongly resembles the *O. polymorpha* chromosome 6 right subtelomere, as the subtelomeric heterochromatin extends ~50 kb from each chromosome end and contains numerous repressed genes despite synteny loss, suggesting the heterochromatin on the *O. haglerorum* chromosome 1 terminus was recreated following repair of the putative single DNA break (Figure 7B, compare the middle two IGV images).

**Figure 7.**
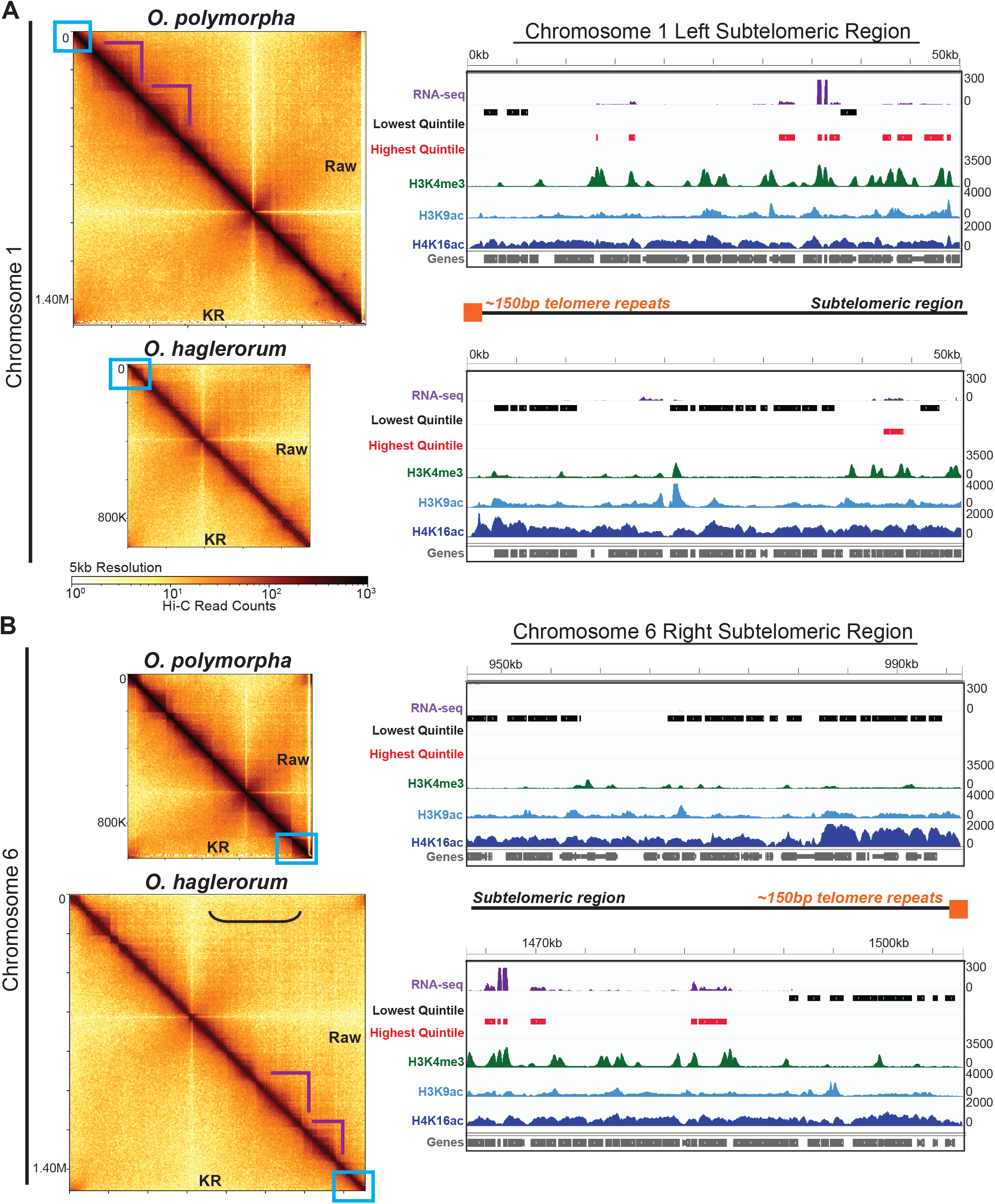
The *O. haglerorum* translocated chromosomes have altered chromatin profiles, relative to the same chromosomes in *O. polymorpha*. (A) Hi-C contact probability heatmaps, as in Figure 5B and scaled to the relative chromosome size, of chromosomes 1 (top two heatmaps) and 6 (bottom two heatmaps) from *O. polymorpha* (top heatmap within each pair) or *O. haglerorum* (bottom heatmap within each pair). Scalebars of log_10_ read counts are shown below the heatmap pairs. Purple triangles show TAD-like structures that may be conserved, and the blue boxes mark the subtelomeric regions displayed in panel B. The bracket in *O. haglerorum* chromosome 6 indicates the region in which intra-chromosomal subtelomeric contacts could occur if chromosome-internal telomere repeats were present (see text). (B) IGV images of ChIP-seq histone PTM enrichment and RNA-seq gene expression levels, as in Figure 3C, of the indicated chromosome 1 (top pair) or chromosome 6 (bottom pair) subtelomeres in *O. polymorpha* (top image within each pair) or *O. haglerorum* (bottom heatmap within each pair).

However, clear differences in the *Ogataea* subtelomeric chromatin exist, particularly when comparing the chromosome pairs. The newly formed *O. haglerorum* chromosome 1 left subtelomere has little similarity to the same subtelomere in *O. polymorpha*, as the *O. polymorpha* heterochromatin extends only ~10 kb from the telomere repeats, while the *O. haglerorum* heterochromatin extends ~40 kb from its chromosome end (Figure 7B, compare the top two IGV images). Further, the newly formed *O. haglerorum* chromosome 6 right subtelomere does not have the same chromatin pattern as its *O. polymorpha* counterpart, as the translocation would have abolished the previously established subtelomeric heterochromatin (Figures 7B [compare the bottom two IGV images], S14). Together, these data suggest that large genome rearrangements have the potential to alter local chromatin patterns in donor and acceptor chromosomes, and that the possibility exists that subtelomeric chromatin within the translocated DNA placed onto the acceptor chromosome is maintained while feature(s) on the donor chromosome are reestablished to preserve chromosome function.

## Discussion

In this work, we report a new, more contiguous *Ogataea haglerorum* genome sequence and leverage this reference genome to decipher changes to the chromatin composition and genome organization between *O. haglerorum* and the closely related species *O. polymorpha*. Our new *O. haglerorum* genome contains seven nearly complete chromosomes, which confirms the overall high level of sequence and synteny conservation with *O. polymorpha* (Hanson et al. 2021). We also show that despite near-contiguous conservation between the two *Ogataea* genomes, there are notable syntenic differences, including the more divergent subtelomeres in each *Ogataea* species, similar to the situation in *Saccharomyces* species (Yue et al. 2017). In fact, the *Ogataea* subtelomeres were previously observed to have a higher SNP density (Hanson et al. 2021). Thus, it is possible that the subtelomeres within the genomes of Saccharomycotina fungi reflect sites of enhanced diversity or continual evolution, relative to chromosomal centers. Comparisons of the synteny between additional fungal genomes within the Ascomycota and the Basidiomycota could expand the conclusions of this hypothesis.

Our data also show that chromatin composition over individual genes is broadly conserved between *O. polymorpha* and *O*. hagleroum. Specifically, in both species, H3K4me3 peaks occur over TSSs, while H3K9 and H4K16 acetylation is primarily enriched over gene bodies. These features are consistent with other ChIP-seq enrichment datasets in the fungi *S. cerevisiae, S. pombe*, and *N. crassa* (Sinha et al. 2006; Shanle et al. 2015; Scadden et al. 2023; Mumford et al. 2024; Zencir et al. 2025), although H3K9ac has been reported to be mainly enriched at yeast gene promoters (Pokholok et al. 2005; Price et al. 2019). The fact that the global histone PTM enrichment patterns over *Ogataea* genes generally mirrors that in other fungi supports the hypothesis that *O. polymorpha* and *O. haglerorum* employ conserved mechanisms for histone PTM deposition to regulate genome function. Specifically, H3K4me3 may act as a molecular memory of RNA Polymerase II-directed transcriptional activity, similar as in *S. cerevisiae* (Ng et al. 2003), while histone acetylation presumably opens chromatin structure (Grunstein 1997; Kuo and Allis 1998; Clayton et al. 2006; Ferraro et al. 2021).

Interestingly, all three histone PTMs are decreased across centromeres and over the silent mating-type genes, while H3K4me3 and H3K9ac enrichment is reduced over subtelomeric genes, suggesting these loci are heterochromatic in both *Ogataea* species. Interestingly, retroelement genes in several heterochromatic centromeres are still transcribed in *O. polymorpha*, as polyadenylated transcripts map to these loci. LTR retrotransposons are thought to have played an important role in centromere evolution in budding yeasts following loss of RNAi-dependent centromere establishment and maintenance, in addition to the loss of the H3K9 methylation mechanism of centromeric heterochromatin formation (Haase et al. 2026). In *Ogataea*, this retroelement-dependent centromeric heterochromatin does not extend into pericentromeric regions, consistent with a lack of pericentromeric heterochromatin in *Candida lusitaniae* (Kapoor et al. 2015), which may be correlated with the different mechanisms budding yeast employ for heterochromatin regulation (Hanson and Wolfe 2017). Subtelomeric chromatin is also silent, as the genes in subtelomeric regions are transcriptionally repressed in *O. polymorpha* and *O. haglerorum*, consistent with heterochromatic subtelomeres being a predominant feature of fungal genomes (Jamieson et al. 2013). Interestingly, we show that subtelomeric genes display hyperacetylated H4K16 in both *Ogataea* species. This hyperacetylation may be required for silent heterochromatin, as the Silent Information Repressor-3 (Sir3) protein is recruited by H4K16ac for subtelomeric repression in *S. cerevisiae* (Oppikofer et al. 2011). In fact, while heterochromatin domains in C. albicans typically have reduced H3K4me3 and hypoacetylated histones, H3K4 and H4K16 can remain acetylated over some subtelomeric genes (Price et al. 2019), suggesting budding yeast employ non-uniform chromatin composition patterns. If these data are applied to the *Ogataea* clade, it suggests that *O. polymorpha* and *O. haglerorum* HDACs could respond to the acetylation state of heterochromatic histones to silence chromatin, yet the underlying mechanism for region-specific HDAC targeting is unknown. One possible mechanism is that HDACs, such as Sir2, form condensed heterochromatin domains refractory to the transcription machinery in these species (Rusche et al. 2003; Hickman et al. 2011; Gartenberg and Smith 2016). NAD^+^ HDAC genes have expanded in the genomes of other yeast species that lost the H3K9me3-catalyzing heterochromatin machinery, including *S. cerevisiae* and *Candida albicans* (Hickman et al. 2011; Rupert et al. 2016), highlighting the reliance on HDAC-dependent heterochromatin silencing in these yeasts. However, a similar expansion of NAD^+^ HDAC genes has not occurred in the *Ogataea* clade, which coupled with the lack of H3K9me3 machinery in these yeasts, means that the mechanism(s) for heterochromatin formation remain unclear. Future experiments characterizing HDAC activity, targets, and regulation in *Ogataea* species will provide insights into the heterochromatin formation mechanism(s).

We also showed that the *O. polymorpha* and *O. haglerorum* genomes organize into the Rabl chromosome conformation, which is highly conserved among other Ascomycete fungi (Duan et al. 2010; Mizuguchi et al. 2014; Galazka et al. 2016; Seidl et al. 2020; Rodriguez et al. 2022; Torres, Reckard, et al. 2023). The Rabl conformation is characterized by a centromere cluster independent of telomere bundles, each of which associates with the nuclear periphery. While our Hi-C data cannot distinguish the number of centromeric or telomeric foci or the subnuclear cluster localization, since our Hi-C data resembles the Hi-C data of organisms with subnuclear microscopic data, including *S. cerevisiae, S. pombe*, and *N. crassa*, these heterochromatic clusters likely localize to the nuclear periphery. Further, considering that the 32 telomeres of the 16 *S. cerevisiae* chromosomes reportedly form three to ten foci in exponentially growing haploid cells, but the 14 telomeres of the seven *Neurospora* chromosomes cluster into two to four foci (Gotta et al. 1996; Hediger et al. 2002; Bystricky et al. 2005; Klocko et al. 2016; Laporte et al. 2016; Ruault et al. 2021), the 14 telomeres of the seven *O. polymorpha* and *O. haglerorum* chromosomes most likely aggregate into two to four foci. Importantly, we found that both the centromeres and telomeres in these species are heterochromatic, consistent with fungal chromatin profiles. Therefore, the localization of these chromosome features to the nuclear periphery, which is known to be a repressive environment in eukaryotes (Taddei et al. 2004; Loïodice et al. 2021; Marin et al. 2025), would promote centromeric and telomeric DNA silencing. Currently, a comprehensive understanding of the factors mediating centromere or telomere clustering, or their nuclear periphery association, are unknown but clues from *S. cerevisiae* and *S. pombe* exist. At the telomeres, the yeast Shelterin complex protects chromosome ends and mediates telomere clustering (Hediger et al. 2006; Kaizer et al. 2015; Steinberg-Neifach and Lue 2015; Wang et al. 2016; Xue et al. 2017; Torres, Reckard, et al. 2023), but the Shelterin complex has not been examined in either *Ogataea* species. Moreover, the proteins mediating interphase centromere clustering are unknown but presumably the centromere-specific histone variant Cse4/CENP-A/CenH3 plays a role (Meluh and Koshland 1997; Camahort et al. 2009; Smith et al. 2012; Seidl et al. 2020; Polisetty et al. 2025). Our Hi-C data also shows that euchromatin in *O. polymorpha* and *O. haglerorum* harbors nested TAD-like structures that may be delineated by highly expressed genes and H3K9ac peaks, implying the euchromatin within chromosome arms could mediate these hierarchical chromatin loop structures. It is possible that the transcription or histone acetylation machinery at TAD-like structure boundaries restrict inter-TAD interactions, or that the TAD-like structures observed here are analogous to the less prominent TAD-like loops regulating replication timing in budding yeasts (Eser et al. 2017). In contrast, filamentous fungi may employ alternative mechanisms to form TAD-like structures, as the TAD-like structure boundaries in *Verticillium dahliae* have reduced gene expression and greater nucleotide diversity (Torres, Kramer, et al. 2023), while TAD-like chromatin loops in *N. crassa* form when two H3K9me3-marked interspersed constitutive heterochromatic regions associate (Rodriguez et al. 2022). It remains possible that cohesin (e.g., the Structural Maintenance of Chromosomes [SMC] complex), which extrudes chromatin loops through a central ring structure and forms off-diagonal chromatin globules in Hi-C experiments (Mizuguchi et al. 2014; Rao et al. 2017; Dauban et al. 2020; Bauer et al. 2021; Davidson and Peters 2021; Zhao et al. 2025), mediates nested loop formation in *O. polymorpha* and *O. haglerorum*. Further work exploring the proteins that form chromatin loops in these species is necessary.

The most striking difference between the *O. polymorpha* and *O. haglerorum* genomes is the translocation between *O. haglerorum* chromosomes 1 and 6 impacting genome organization and chromatin composition. Specifically, the TAD-like structures in chromosomes involved in the translocation, as well as in unaffected syntenic chromosomes, appear similar in both species, suggesting conserved mechanisms to establish and maintain these nested, modular TAD-like structures. However, we hesitate to make definitive conclusions about any inter-species TAD-like structure conservation given the challenges in predicting these structures, since chromatin loops in fungi can be quite heterogeneous (Rodriguez et al. 2022). Importantly, the formation of required chromosomal features must be paramount to any chromatin changes. For example, telomeric repeats on each chromosome must be present following any translocation. We show the *O. haglerorum* chromosome 1 left arm telomere is present after translocation repair, which is necessary for silencing the newly positioned subtelomeric genes. In contrast, the telomere placed internally in the *O. haglerorum* acceptor chromosome 6 following translocation repair has not been retained, as no chromosome-internal intra- or inter-telomere contacts are observed, suggesting any remaining telomere sequence relict(s) in the chromosome center were lost following translocation repair. Indeed, ectopic, chromosome-internal telomere repeats can three-dimensionally associate with telomere repeats on chromosome ends in *Neurospora* (Mumford et al. 2026), so had telomere repeats been retained in *O. haglerorum* chromosome 6, long-range inter-telomere repeat interactions would have been obvious. We note that it is unclear if this *O. haglerorum* translocation was a quasi-terminal genome rearrangement, in which two double stranded DNA breaks caused the telomere repeats to remain on chromosome 1, or if the telomere repeats on chromosome 1 were added post-translocation. Regardless, the need for fungi to protect the linear DNA on chromosome ends as well as cluster subtelomeres is paramount for genome function; our evidence suggests that subtelomeric heterochromatin was historically reestablished post-translocation. Overall, the conservation of subtelomeres between *O. polymorpha* and *O. haglerorum* suggests that modular, sequence dependent and independent features exist at chromosome ends. We propose that stable genome synteny changes involving chromosome termini must establish telomeric repeat DNA for end protection and (sub)telomere clustering.

Much remains unknown about how genome organization or chromatin composition can drive speciation. We show that in addition to the well-established nucleotide differences between yeast species, species-specific chromatin changes can locally occur despite the broad conservation of genome function. Our work also highlights how large genome rearrangements can alter histone PTM deposition and transcription patterns. While the translocation observed in the *O. haglerorum* strain studied here is not conserved among other *O. haglerorum* isolates, multiple other large-scale rearrangements have sporadically occurred throughout the *O. polymorpha* species complex (Hanson et al. 2021). We speculate that translocation-driven chromatin changes may play a role in speciation or niche survivability; at the very least, translocations in these fungi are reasonably tolerated, highlighting the plasticity of fungal genomes. We suggest that alterations to any component of genome function, including genome organization, chromatin composition, and gene expression – which may be driven by DNA sequence changes – may facilitate species divergence. While our work has helped clarify how genome function changes may relate to diversification, future research could elucidate how genome function changes may facilitate speciation in other species, including metazoans.

## Supporting information

Supplemental_File_S2_fasta

Supplemental_File_S3_gff

## Data Availability

Supplemental Figures, Files, and Tables are available on the GSA Figshare portal. File S1 contains detailed Materials and Methods, and Files S2 and S3 are the fasta and gff files for the refined *O. haglerorum* reference genome. Table S1 reports conserved histone deacetylase genes in the genomes of *Ogataea* species, while Table S2 reports genome statistics for the *O. polymorpha* and *O. haglerorum* genomes. Oxford Nanopore Technology long read whole genome sequencing data was deposited to the NIH NCBI Sequence Read Archive (SRA) under accession number SRR37174755. The *O. haglerorum* reference genome is available at the NCBI Genome database (Accession number JBZEMS000000000) and the fasta (File S2) and gff (File S3) are available at FigShare associated with this manuscript. All RNA-seq, ChIP-seq, and Hi-C data was deposited to the NCBI GEO under superseries accession number GSE318849, with individual accession numbers for ChIP-seq (GSE318721), RNA-seq (GSE318843), and Hi-C (GSE318848) datasets.

## Acknowledgements

The authors would like to thank Hanson lab members Mai Nguyen and Collin Ralston, and all members of the Klocko lab, as well as members of the Colorado College Department of Molecular Biology and the UCCS Departments of Chemistry & Biochemistry and Biology for helpful discussions regarding this project. The authors also thank *G3* editor Minou Nowrousian and the three anonymous reviewers whose feedback helped improve this manuscript. This project was funded by start-up funds from the UCCS College of Letters, Arts, and Sciences to AD Klocko and by the Dean of the Faculty’s Office and Natural Sciences Division of Colorado College to SJ Hanson. In addition, TJ Lundberg, NM Lande, and AD Klocko were supported by NIH grant R15GM140396 during this project and NIH grant R35GM163528 during the final stages of peer review.

