## Supplemental_Figure_S1 for "A Translocation within the *Ogataea* Species Complex Alters Local Subtelomeric Chromatin while Maintaining Overall Genome Organization"

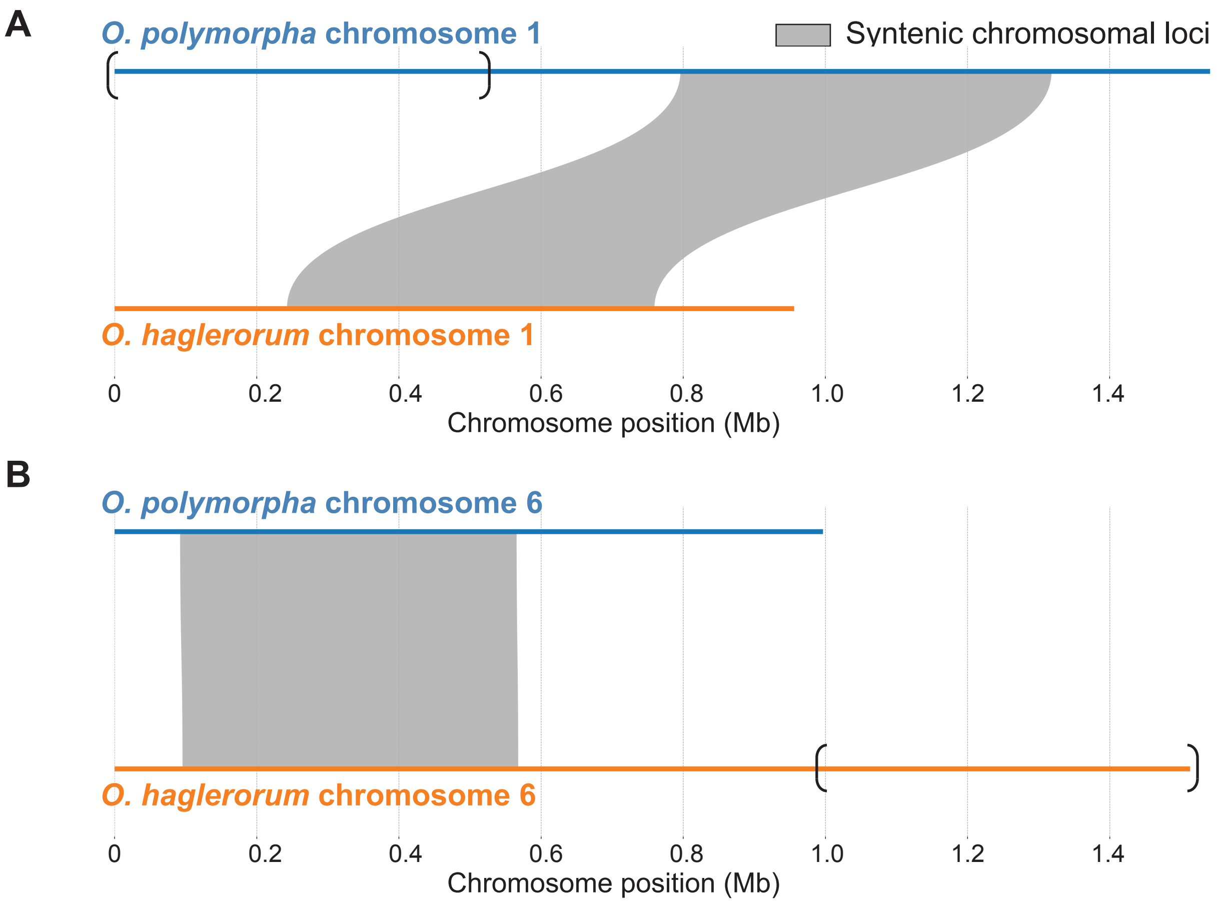


**Figure S1. The two chromosomes from two *Ogataea* species involved in the translocation have syntenic regions when compared.** (A-B) Synteny plots showing the sequence conservation between *O. polymorpha* chromosomes (blue lines) and *O. haglerorum* (orange lines) for chromosomes (A) 1 or (B) 6. Gray shading shows syntenic loci. Brackets show the section of *O. polymorpha* chromosome 1 that is translocated onto *O. haglerorum* chromosome 6, as previously reported (Hanson et al. 2021) and confirmed by Hi-C (this work).
