## Supplemental_Figure_S2 for "A Translocation within the *Ogataea* Species Complex Alters Local Subtelomeric Chromatin while Maintaining Overall Genome Organization"

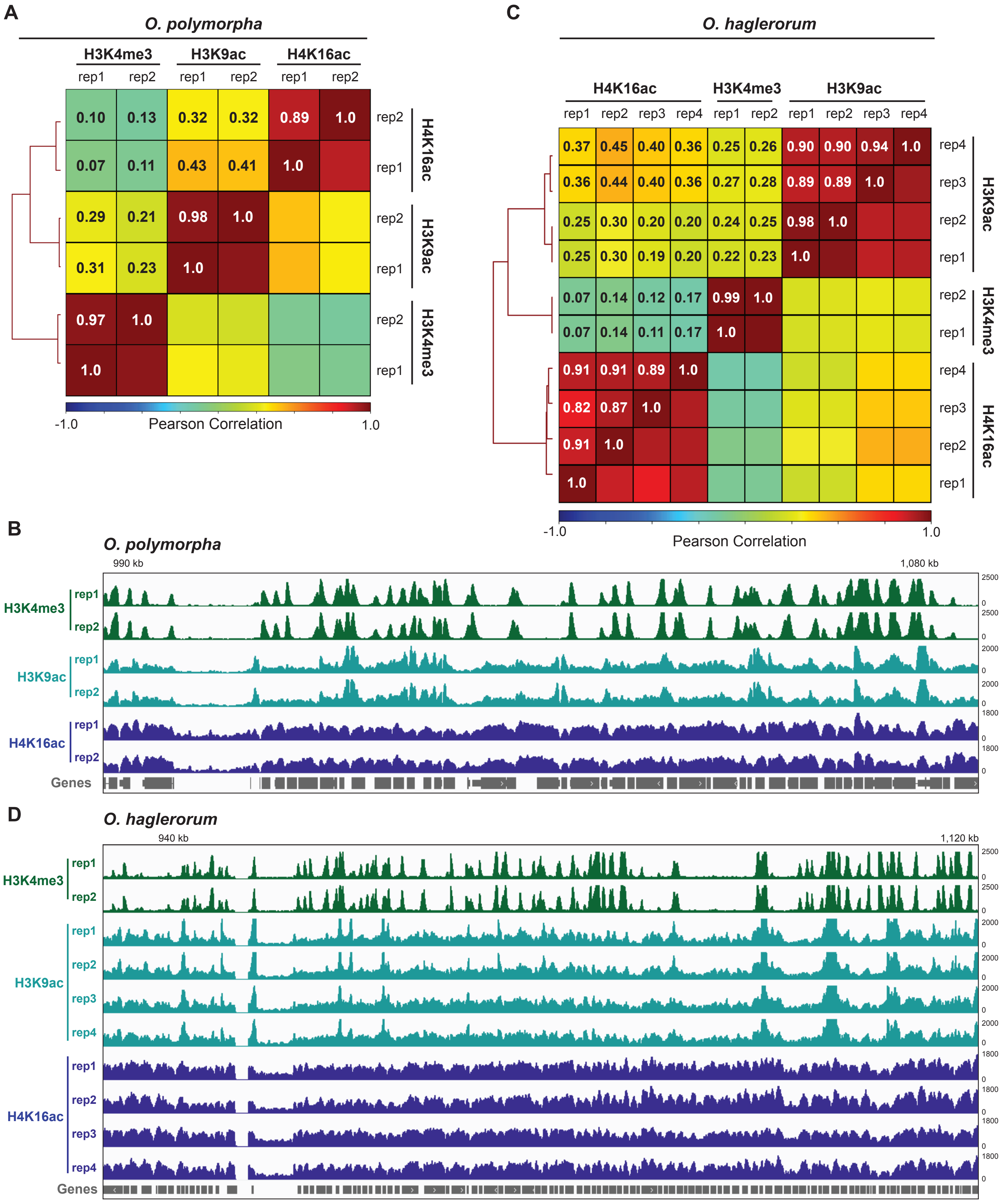


**Figure S2. All Chromatin Immunoprecipitation-sequencing (ChIP-seq) experiments from *Ogataea* species are reproducible.** (A,C) Heatmaps of the Pearson correlation values and IGV images of bigwig tracks (B,D), colored as in Figure 1, of the H3K4me3, H3K9ac, and H4K16ac ChIP-seq experiments from *O. polymorpha* (A-B) or *O. haglerorum* (C-D). ChIP-seq replicate numbers are indicated in each panel.
