## Supplemental_Figure_S3 for "A Translocation within the *Ogataea* Species Complex Alters Local Subtelomeric Chromatin while Maintaining Overall Genome Organization"

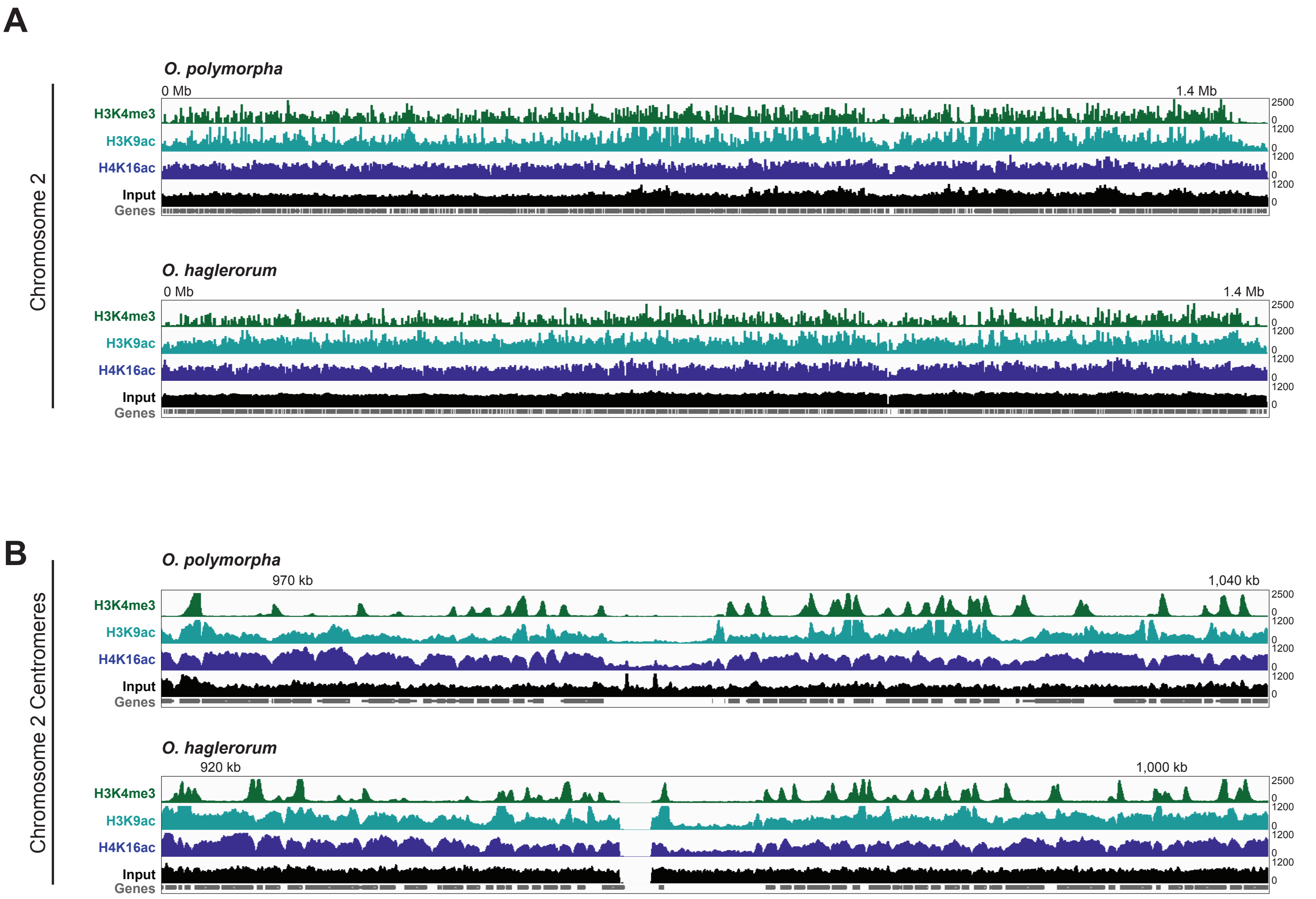


**Figure S3. "ChIP-seq experiments for histone PTMs in *O. polymorpha* and *O. haglerorum* genomes are specific when compared to input DNA sequenced files.** IGV images displaying the H3K4me3, H3K9ac, and H4K16ac ChIP-seq enrichment, plotted as in Figure S4, or the input DNA that went into the ChIP-seq experiments (black) across (A) chromosome 2 or the (B) chromosome 2 centromere of *O. polymorpha* (top) or *O. haglerorum* (bottom)
