## Supplementary figures and images for "A Translocation within the *Ogataea* Species Complex Alters Local Subtelomeric Chromatin while Maintaining Overall Genome Organization"

### Supplemental_Figure_S4Supplemental_Figure_S3

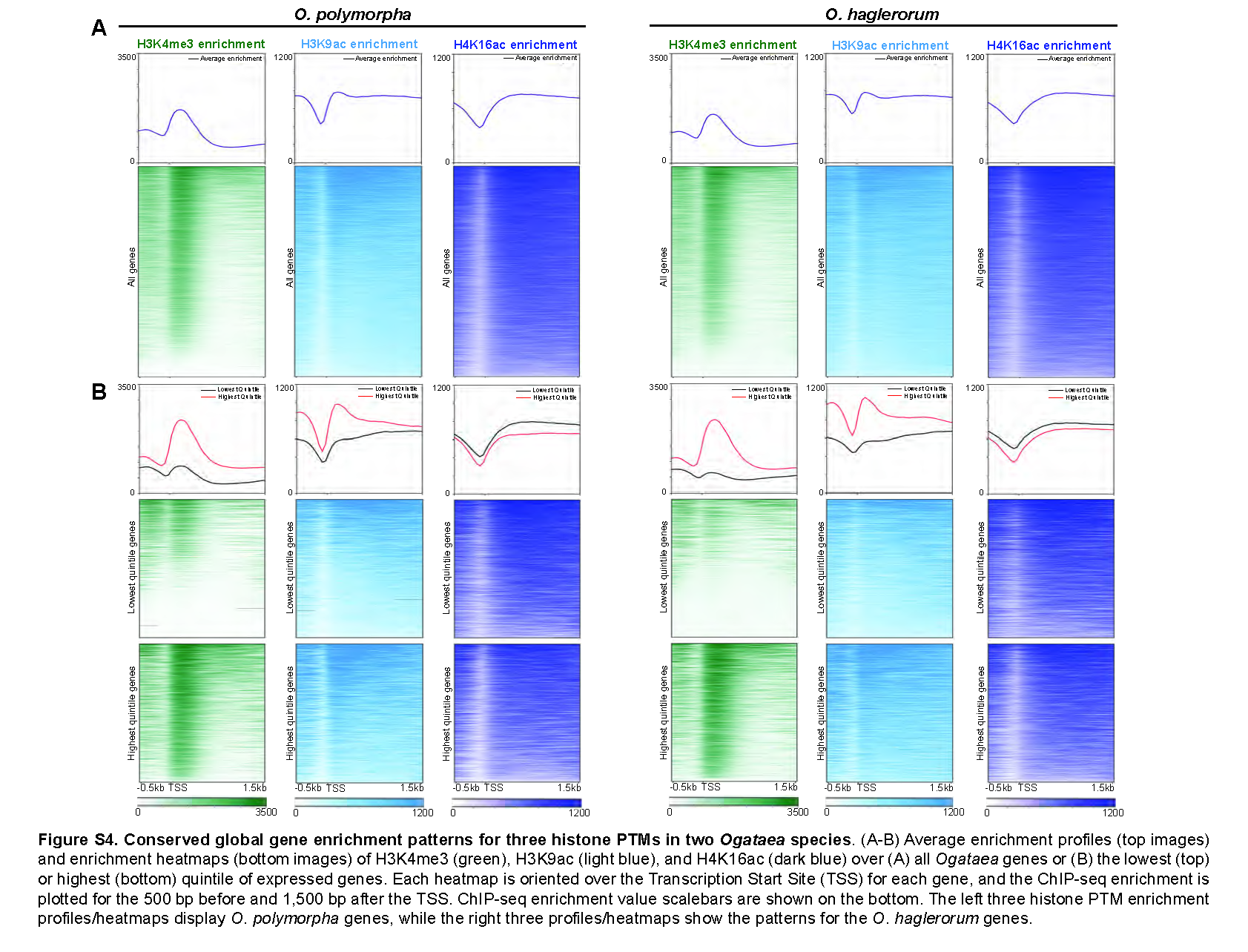
