## Supplemental_Figure_S5 for "A Translocation within the *Ogataea* Species Complex Alters Local Subtelomeric Chromatin while Maintaining Overall Genome Organization"

### *O. polymorpha* Centromeres

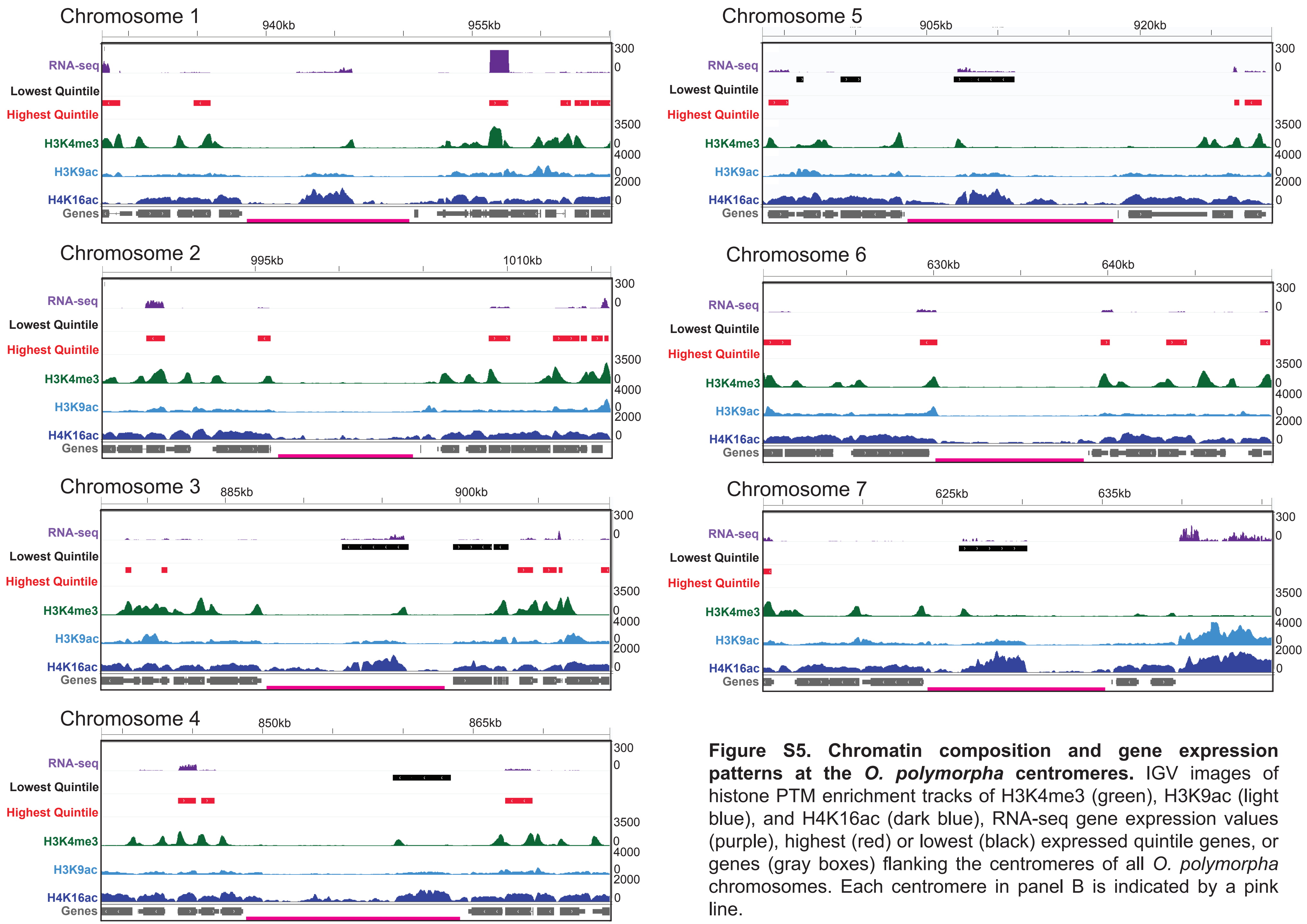


**Figure S5. Chromatin composition and gene expression patterns at the *O. polymorpha* centromeres.** IGV images of histone PTM enrichment tracks of H3K4me3 (green), H3K9ac (light blue), and H4K16ac (dark blue), RNA-seq gene expression values (purple), highest (red) or lowest (black) expressed quintile genes, or genes (gray boxes) flanking the centromeres of all *O. polymorpha* chromosomes. Each centromere in panel B is indicated by a pink line.
