## Supplemental_Figure_S6 for "A Translocation within the *Ogataea* Species Complex Alters Local Subtelomeric Chromatin while Maintaining Overall Genome Organization"

### *O. haglerorum* Centromeres

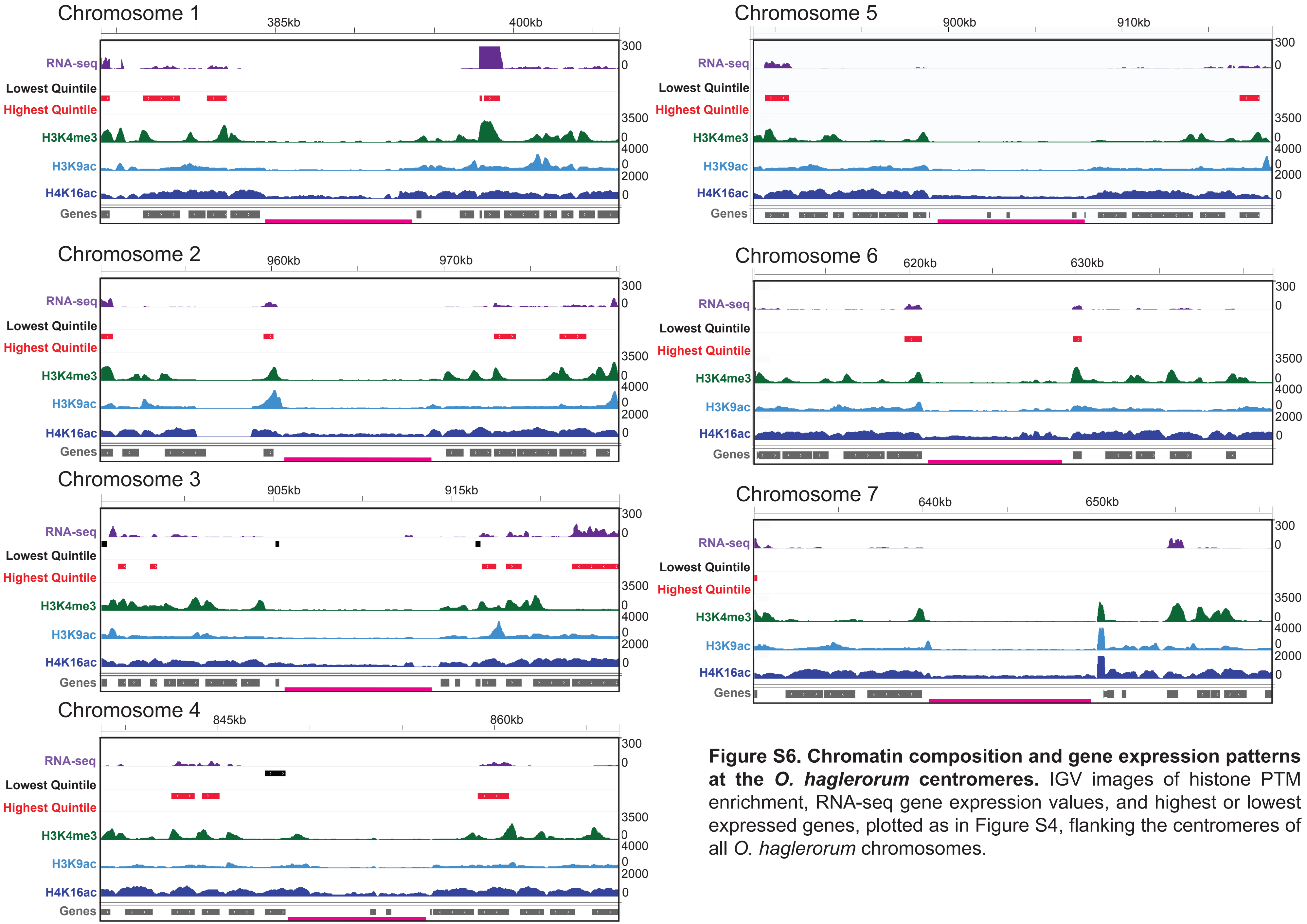


**Figure S6. Chromatin composition and gene expression patterns at the *O. haglerorum* centromeres.** IGV images of histone PTM enrichment, RNA-seq gene expression values, and highest or lowest expressed genes, plotted as in Figure S4, flanking the centromeres of all *O. haglerorum* chromosomes.
