## Supplemental_Figure_S8 for "A Translocation within the *Ogataea* Species Complex Alters Local Subtelomeric Chromatin while Maintaining Overall Genome Organization"

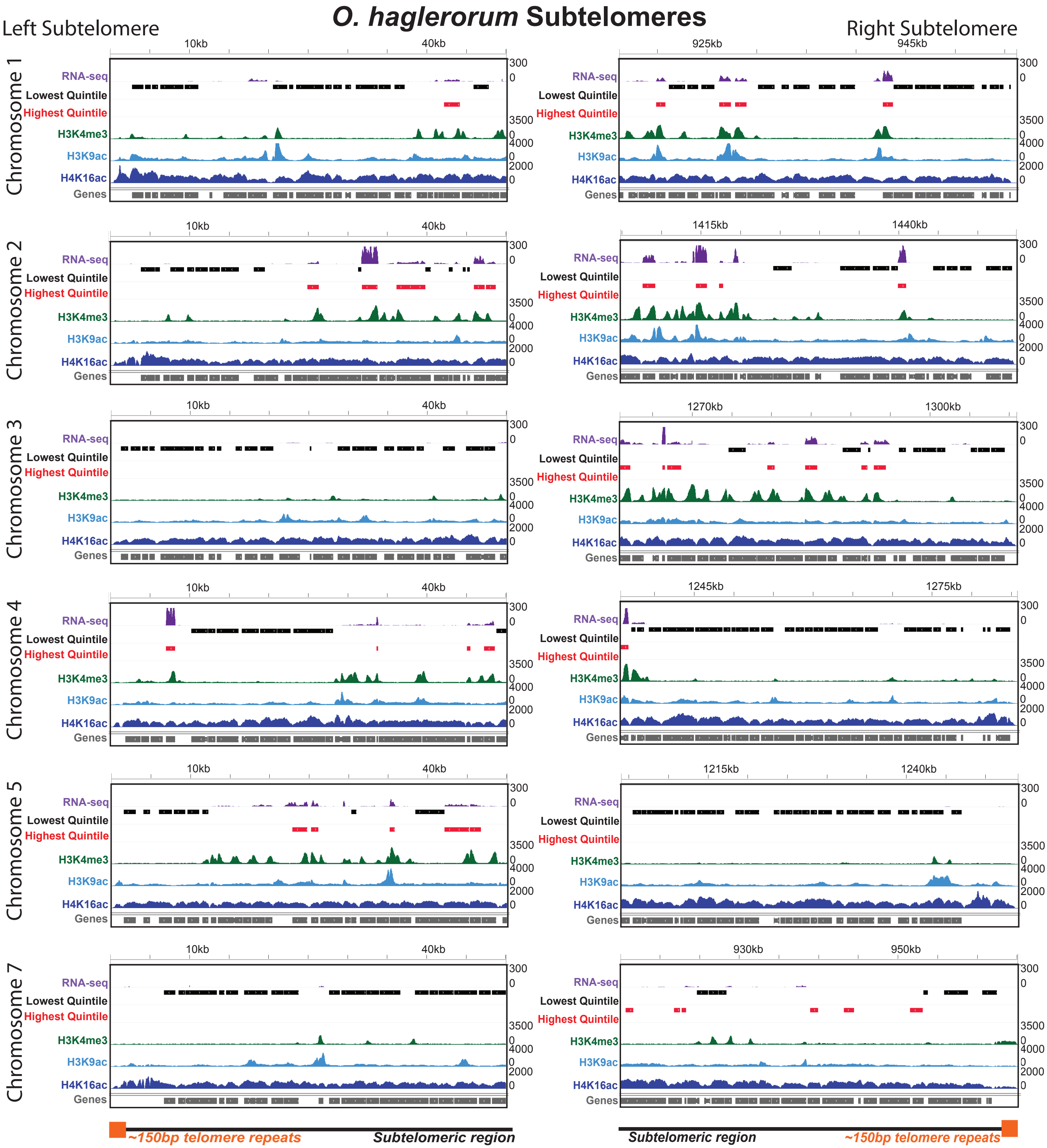


**Figure S8. Chromatin composition and gene expression patterns at the *O. haglerorum* telomeres.** IGV images of histone PTM enrichment, RNA-seq gene expression values, and highest or lowest expressed genes, plotted as in Figure S4, of the chromatin comprising the 50 kb immediately adjacent to the telomeres (the subtelomeric chromatin) of all *O. haglerorum* chromosomes.
