## Supplemental_Figure_S9 for "A Translocation within the *Ogataea* Species Complex Alters Local Subtelomeric Chromatin while Maintaining Overall Genome Organization"

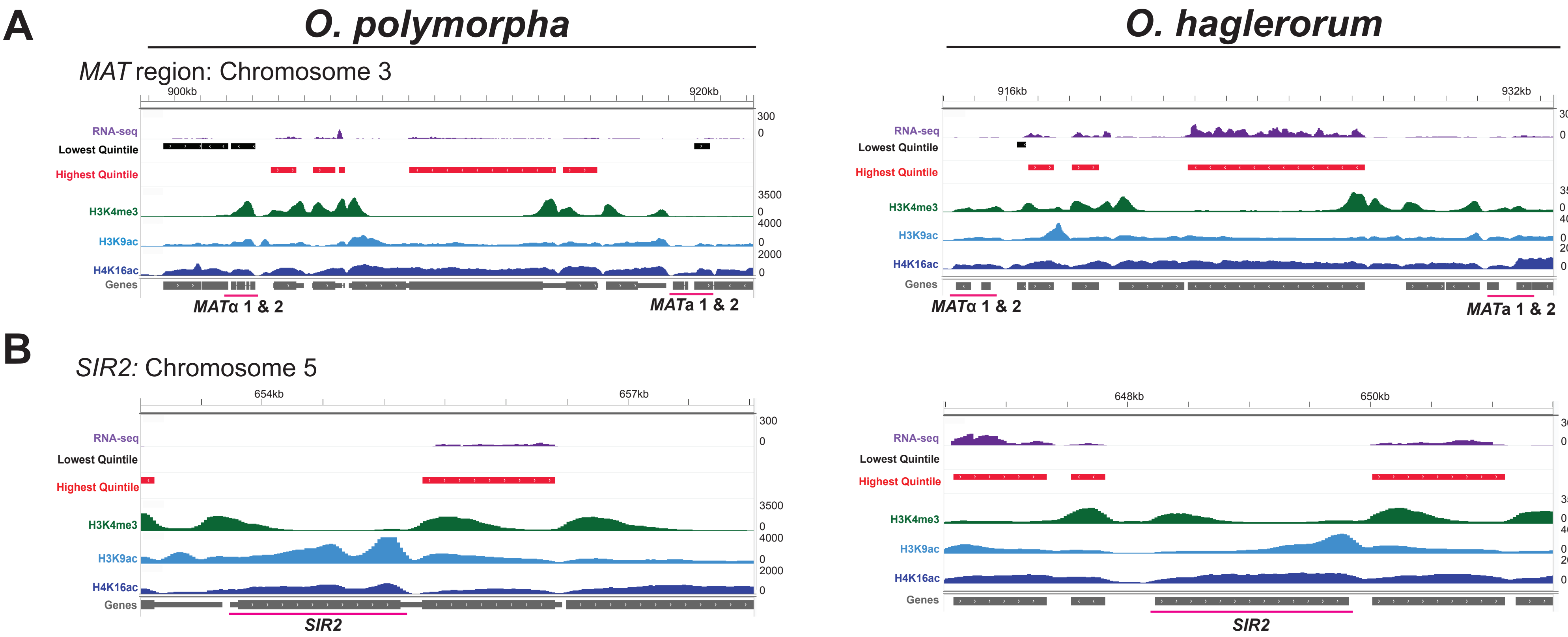


**Figure S9. Distinct expression levels and chromatin composition patterns in *O. polymorpha* and *O. haglerorum* genes.** (A-B) IGV images of histone PTM enrichment and gene expression values, plotted as in Figures 2B-C, over select genes, including (A) the mating type locus (the chromosome 3 centromere is to the left in each image) and (B) the *sir2* histone deacetylase gene.
