## Supplemental_Figure_S10 for "A Translocation within the *Ogataea* Species Complex Alters Local Subtelomeric Chromatin while Maintaining Overall Genome Organization"

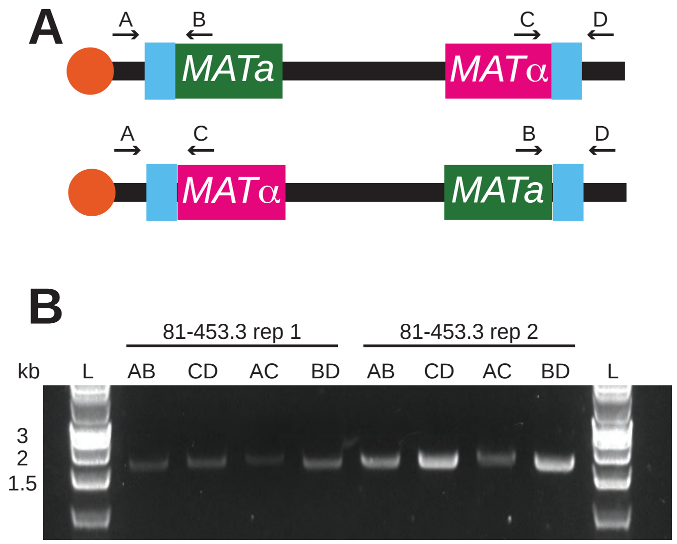


**Figure S10. *O. haglerorum* samples contain a mixture of MATa and MATα mating types.** (A) Diagram showing two possible orientations of the *O. haglerorum* mating type region with primer positions (A-D) shown. The positions of the centromere (orange circles) and inverted repeat sequences (blue boxes) are also indicated. (B) Agarose gel image for mating type region PCR. Ladder lanes (L) and band sizes in kilobases (kb) are indicated. Primer combinations are listed for each lane for each replicate used in ChIP and Hi-C.
