## Supplemental_Figure_S11 for "A Translocation within the *Ogataea* Species Complex Alters Local Subtelomeric Chromatin while Maintaining Overall Genome Organization"

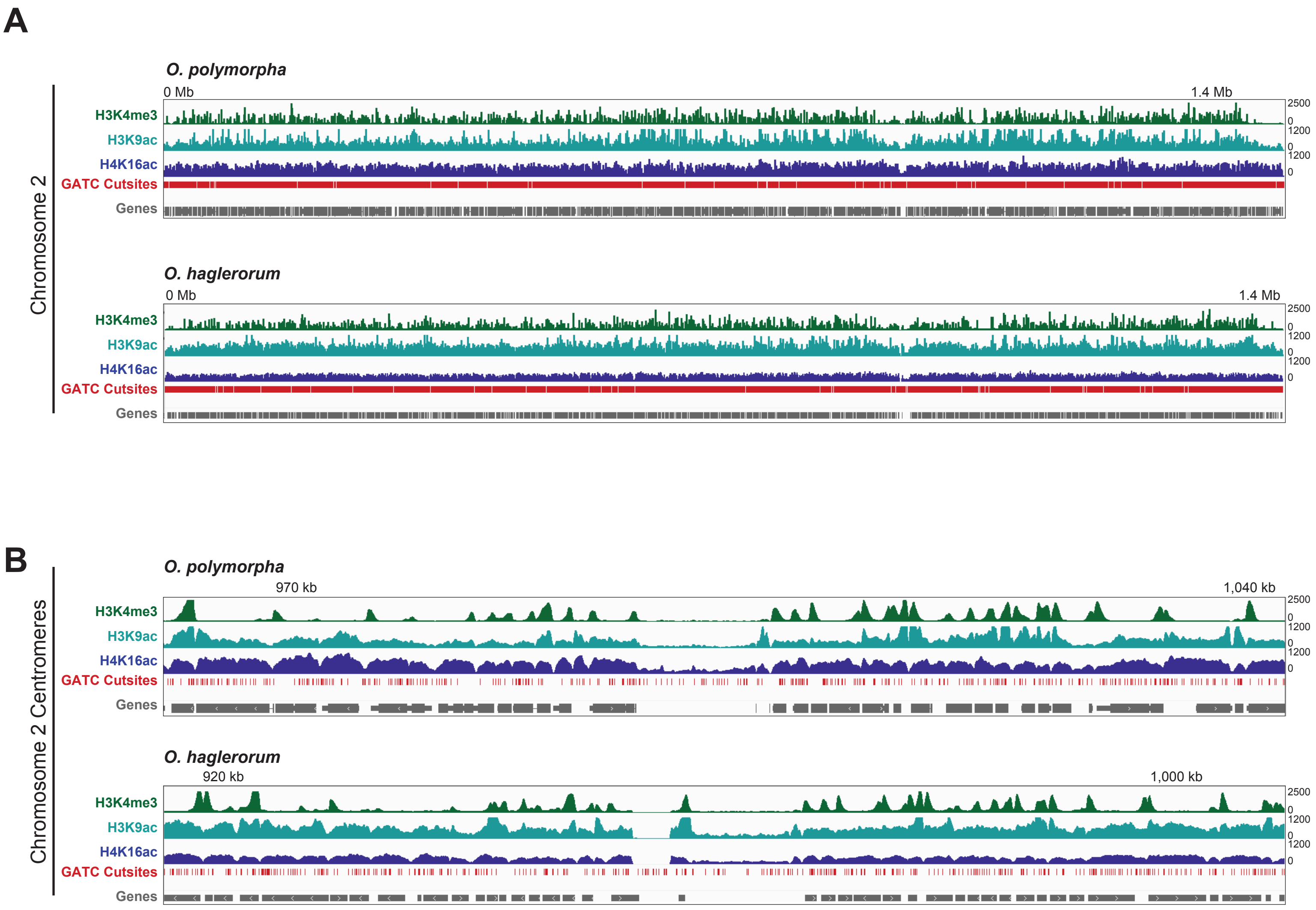


**Figure S11. Recognition sequences for restriction enzyme *Dpn*II have an approximate even distribution across the *O. polymorpha* and *O. haglerorum* genomes.** IGV images displaying the H3K4me3, H3K9ac, and H4K16ac ChIP-seq distribution, plotted as in Figure S4, or the distribution of *Dpn*II sites (5' GATC; red) across (A) chromosome 2 or the (B) chromosome 2 centromere of *O. polymorpha* (top) or *O. haglerorum* (bottom).
