## Supplemental_Figure_S12 for "A Translocation within the *Ogataea* Species Complex Alters Local Subtelomeric Chromatin while Maintaining Overall Genome Organization"

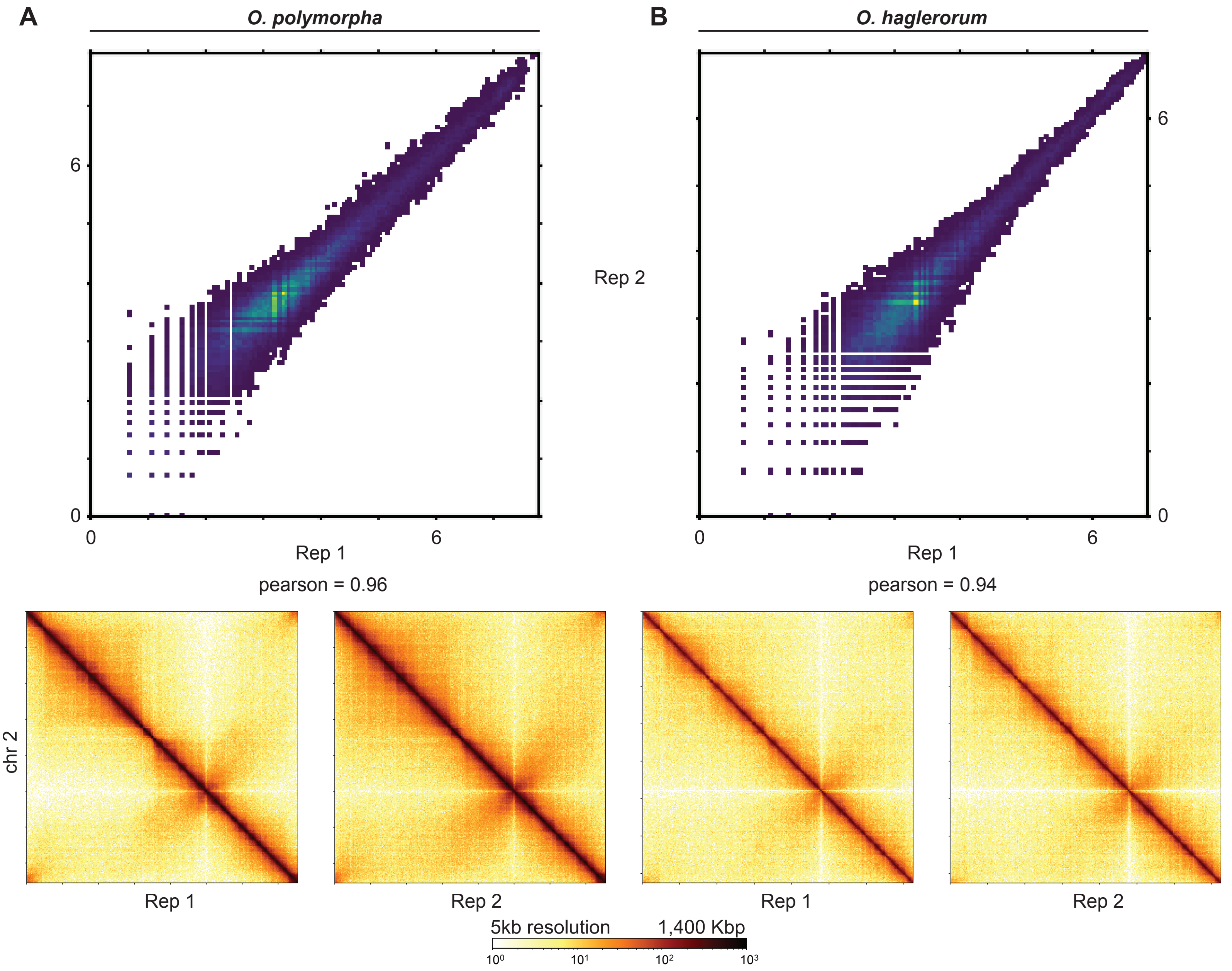


**Figure S12. Replicate Hi-C experiments in either *Ogataea* species are well correlated.** (A-B) Scatterplots comparing the log1p values of the two Hi-C replicates (top) or heatmaps displaying the contact probability across chromosome 2 of each replicate (bottom) for (A) *O. polymorpha* or (B) *O. haglerorum*. Pearson correlation values between the replicates are reported below the scatterplots.
