## Supplemental_Figure_S13 for "A Translocation within the *Ogataea* Species Complex Alters Local Subtelomeric Chromatin while Maintaining Overall Genome Organization"

### *O. haglerorum* Hi-C Mapped to *O. polymorpha* Genome

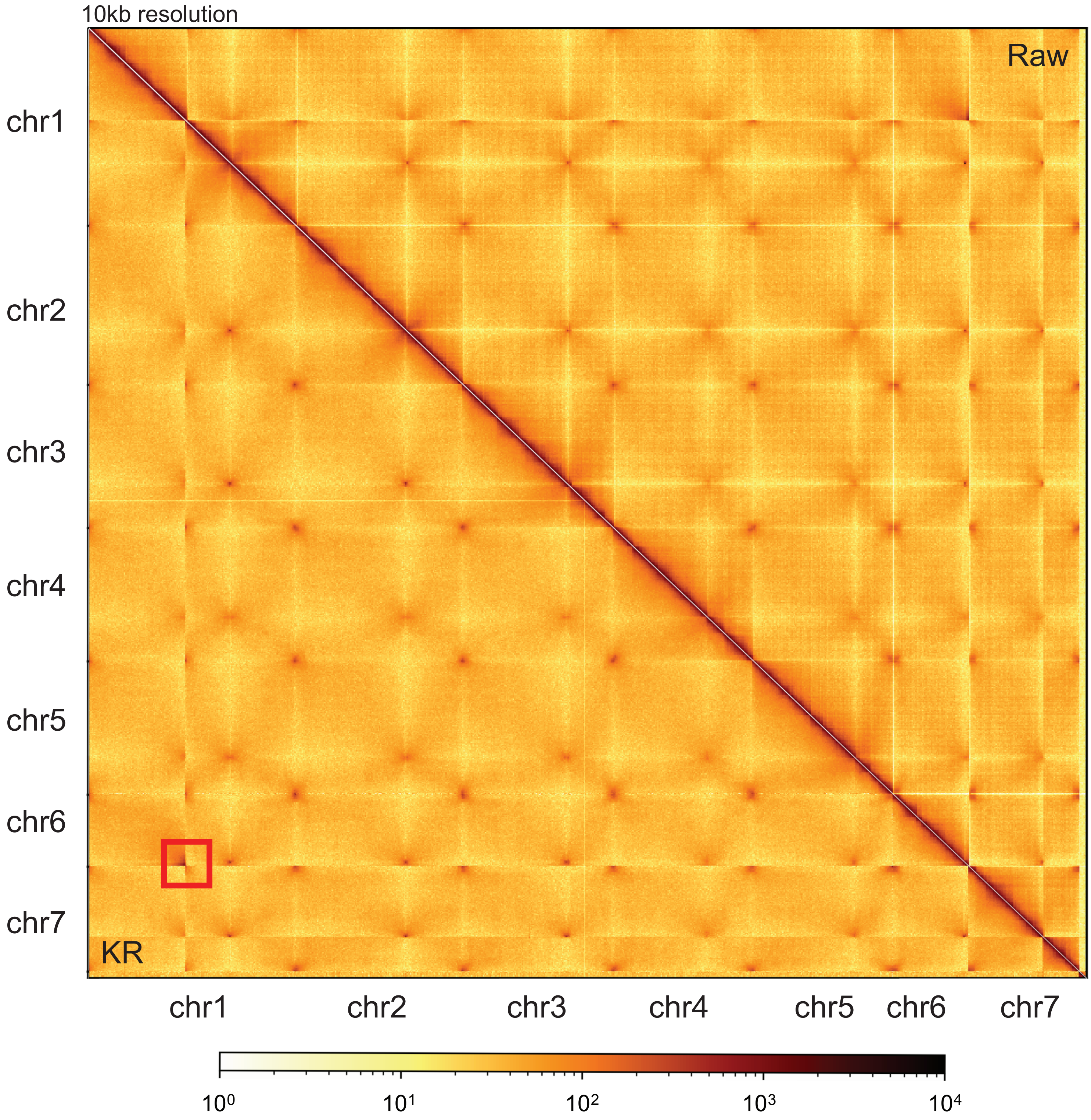


**Figure S13. Hi-C confirms the large genome rearrangement between chromosomes 1 and 6 in *O. haglerorum,* relative to *O. polymorpha* genome synteny.** Hi-C contact probability heatmaps of the whole *O. haglerorum* genome at 10 kb resolution. Hi-C data was mapped to the *O. haglerorum* genome that retained the chromosome synteny of the *O. polymorpha* genome. Raw Hi-C data is shown above the diagonal while the Knight-Ruiz corrected (KR) contact probability heatmap to reduce underlying sequencing biases, is displayed below the diagonal. A scalebar displaying the log10 read counts is shown below the heatmap. The translocation breakpoint between chromosomes 1 and 6 in *O. haglerorum* is indicated by the red box.
