## Supplemental_Figure_S14 for "A Translocation within the *Ogataea* Species Complex Alters Local Subtelomeric Chromatin while Maintaining Overall Genome Organization"

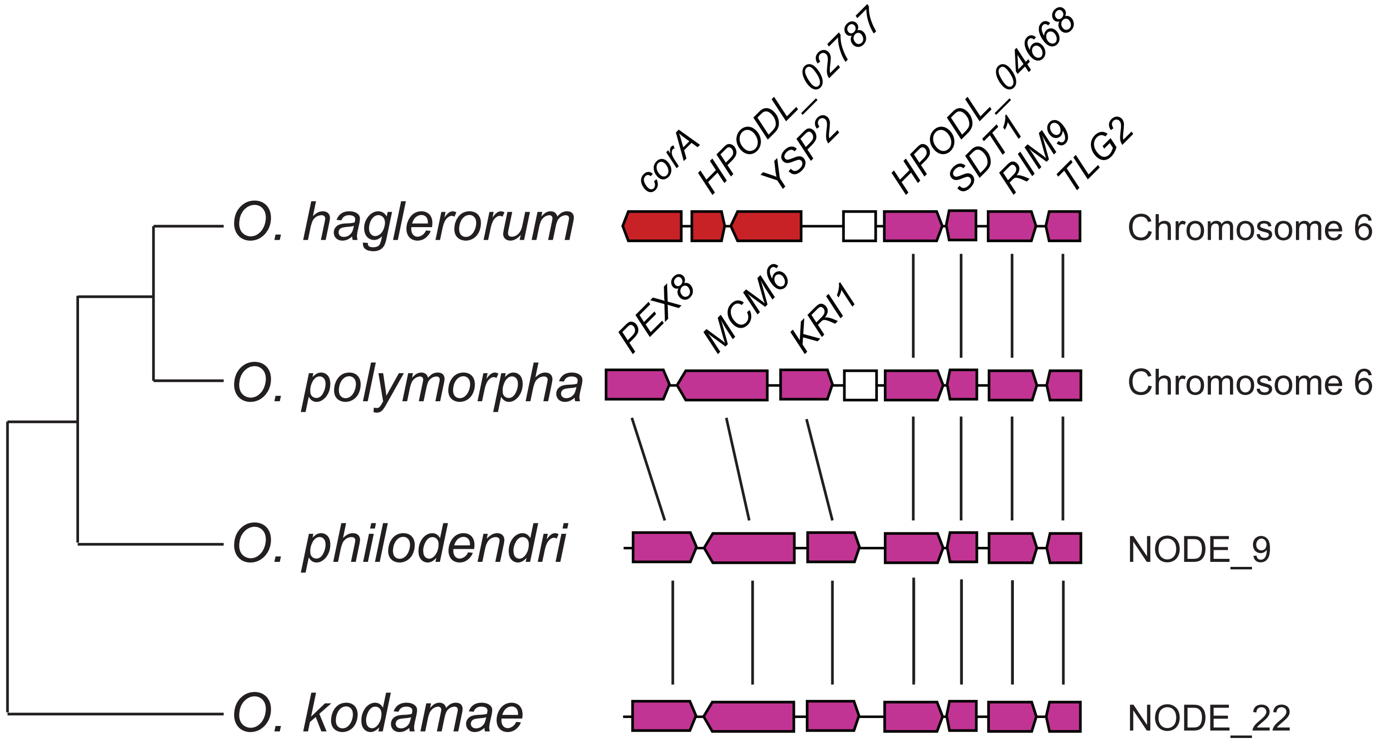


**Figure S14. Synteny comparison in *Ogataea* species.** The translocation identified in *O. haglerorum* strains 81-453.3 and 81-461.3 is not shared in the outgroup species *O. philodendri* (NRRL Y-7210, GCA_003706115.3) or *O. kodamae* (NRRL Y-17234, GCA_003706165.3). Phylogenetic tree is based on (Hanson et al. 2021). Red genes are found on chromosome 1 in *O. polymorpha* and pink genes are found on chromosome 6 in *O. polymorpha*. Vertical lines indicate shared synteny between genomes, with the scaffold/contig names indicated.
