## Supplemental_Figure_S15 for "A Translocation within the *Ogataea* Species Complex Alters Local Subtelomeric Chromatin while Maintaining Overall Genome Organization"

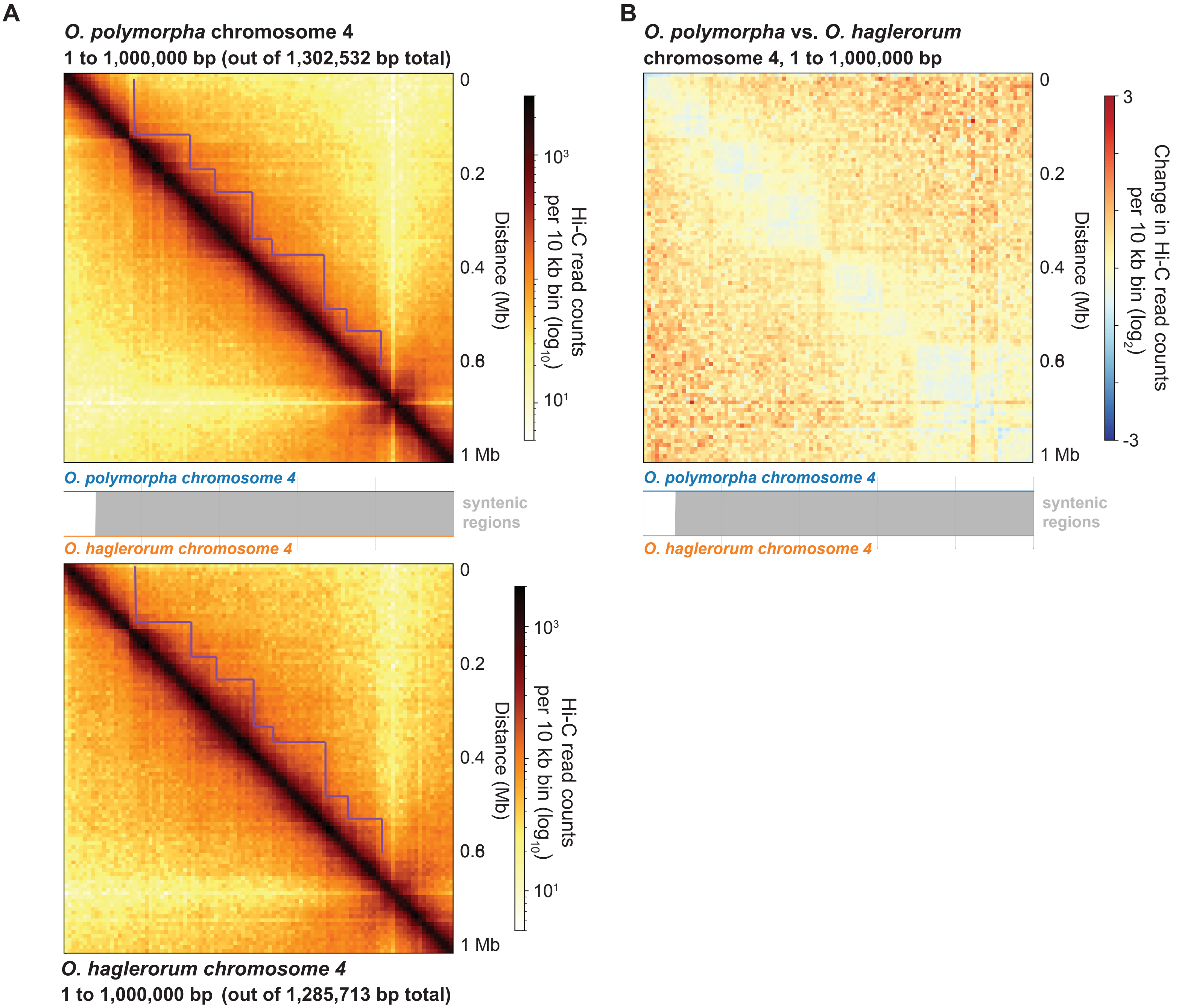


**Figure S15. The primarily syntenic chromosome 4 of *O. polymorpha* and *O. haglerorum* forms similar TAD-like structures in both species.** (A) Hi-C contact probability heatmaps of the first 1,000,000 basepairs, oriented from the left telomere, of chromosome 4 from both *O. polymorpha* (top) and *O. haglerorum* (bottom) at 10 kb resolution. A plot showing chromosome 4 synteny between the two Ogataea species is shown between both heatmaps. The purple line in both heatmaps highlights the similar TAD-like structures found in this syntenic chromosome. (B) Heatmap showing the $log_2$ change in Hi-C contact probability between *O. polymorpha* and *O. haglerorum* across the syntenic region of chromosome 4 displayed in panel A.
