## Supplemental_File_S1 for "A Translocation within the *Ogataea* Species Complex Alters Local Subtelomeric Chromatin while Maintaining Overall Genome Organization"

**File S1 - Supplemental Materials and Methods for:**

### A Translocation within an Ogataea Clade Species Alters Local Subtelomeric Chromatin while Maintaining Global Genome Organization

Tiffany J. Lundberg, Nickolas M. Lande, Daniel Turevski, Riley Figueroa, Sara J. Hanson, Andrew D. Klocko

#### Supplemental Materials and Methods

*Inventory of histone deacetylase genes*

Candidate homologous sequences for histone deacetylases were identified using a Python (version 3.10.11) workflow. An amino acid query sequence was used to identify similar sequences in a specified species based on taxonomic identifications (taxIDs). Using the provided taxIDs, NCBI Datasets and Dataformat (version 15.10.0)(Sayers et al. 2021) were used to gather genome and proteome data to create a species search set. BLAST+ (version 2.14.0)(Altschul et al. 1990) was used to carry out an iterative and reciprocal best hit search process. ETE Toolkits (ete3 version 3.1.1)(Huerta-Cepas et al. 2016) was used to sort species in the protein hit list into their respective family rank to facilitate iterative searches with a related species sequence as a query. The source code for mign is available at Github at https://github.com/maitiennguyen/mign.

*O. haglerorum* MAT *PCR*

The orientation of the mating type locus in *O. haglerorum* was determined by PCR using primers that flank the inverted repeats (A: 5’ – GCCAACAAAGTCATGTGCTCTG – 3’, B: 5’ – GTCGTTGAAGATGTATTTCAGG – 3’, C: 5’ – GGTACGGTGTGCTATTGAATTC – 3’, D: 5’ – GTGCAAGACTTGATCTGGC – 3’). GoTaq Green Master Mix (Promega) was used according to manufacturer’s instructions. PCR was performed for 25 cycles with a 55°C annealing temperature and 1 minute extension. Products were separated on 1% agarose gel with 1X GelRed and 1kb+ ladder (New England Biolabs) for size reference.

*Chromatin Immunoprecipitation-sequencing (ChIP-seq) library preparation*

To prepare *Ogataea* cell pellets for ChIP, overnight cultures grown in YPD broth were backdiluted and grown to log-phase ($OD_{600}$ 0.8-1.2), crosslinked in 1% v/v formaldehyde for 10 minutes at room temperature, quenched with 0.3M glycine for 10 minutes at room temperature, pelleted by centrifugation at 1,800xg and stored at -80°C. Pellets were resuspended in 0.25mL ChIP Lysis buffer (50mM HEPES, pH 7.5; 140 mM NaCl; 1mM EDTA; 1% Triton X-100) with 1x Halt Protease Inhibitor Cocktail (ThermoFisherScientific, cat# 78441). Cells were lysed and chromatin was sheared by a Bioruptor Pico, using a 15-minute protocol of 30 seconds of sonication and 30 second off. Cell debris was pelleted by centrifugation at 5500rpm for 5 minutes, the supernatant was transferred to a 2.0mL screwtop tube, and 2μL of antibody was added and nutated overnight at 4°C. The antibodies used for ChIP-seq were α-H3K4me3 (ActiveMotif cat# 39060, lot# 31420006); α-H3K9ac (ActiveMotif cat# 39137, lot# 28720002), and α-H4K16ac (ActiveMotif cat# 39068, lot# 08719003). The antibody/histone PTM/DNA complex was bound by Protein A/G magnetic beads (washed and resuspended in ChIP Lysis Buffer; ThermoFisherScientific cat# PI78609) for 2 hours at 4°C, and beads were washed twice with ChIP Lysis Buffer, once with ChIP Lysis Buffer + 0.5M NaCl, once with LiCl wash buffer (10mM Tris-HCl pH 8.0; 250mM LiCl; 5% IGEPAL CA-630; 1mM EDTA), and once with TE (10mM Tris-HCl pH 8.0; 1mM EDTA); all washes were for 10 minutes at 4°C. DNA was eluted in TES buffer (50mM Tris-HCl pH 8.0, 10 mM EDTA, 1% SDS), decrosslinked overnight at 65°C, and digested with 100μg/mL RNaseA for 2 hours at 50°C followed with 500μg/mL proteinase K digestion for 2 hours at 50°C. ChIP DNA was organically extracted once with phenol/chloroform/isoamyl alcohol (25:24:1; ThermoFisherScientific cat# AC327115000) and once with chloroform, and ethanol precipitated at -80°C overnight with 2μL of 20mg/mL glycogen (40μg [final]; ThermoFisherScientific cat# R0561) and quantified by Qubit 3.0 fluorimeter using the DNA High Sensitivity (HS) kit.

*Poly-adenine messenger RNA-sequencing (RNA-seq) library preparation*

Overnight YPD cultures were backdiluted and grown to log phase, pelleted and resuspended in TES buffer (10mM Tris HCl, pH7.5, 10mM EDTA pH 8.0, 0.5% SDS). One volume of acid phenol (Invitrogen) was added prior to incubation for 60 minutes at 65°C, with vortexing every 10 minutes. Samples were centrifuged for 5 minutes at 16,000 x g at 4°C, and one additional round of acid phenol and one round of chloroform extraction were performed, followed by ethanol precipitation. Samples were treated with DNase I (Sigma-Aldrich) for 15 minutes at room temperature, followed by incubation with 2mM EDTA at 65°C for 10 minutes. Samples were purified using the RNA Clean and Concentrator-25 kit (Zymo Research) according to manufacturer's instructions. Sample quantity was evaluated with a Qubit 3.0 using the RNA Broad Range kit (ThermoFisher) and integrity was evaluated by TapeStation (Agilent).

*Chromatin Conformation Capture with high-throughput sequencing (Hi-C) library preparation*

Crosslinked Ogataea cell pellets were prepared similarly as ChIP-seq and were stored at -80°C. One pellet was sacrificed to determine the DNA concentration by resuspending the pellet in TE with 0.5% SDS, 4mM EDTA, and 500µg/mL proteinase K and incubating the cells at 65°C overnight, then digesting the pellets with 80µg/mL RNaseA at 37°C for 30 minutes, organically extracting the DNA with phenol/chloroform/isoamyl alcohol (IAA) and chloroform, precipitating the DNA with sodium acetate and isopropanol, resuspending the DNA pellet in 50µL TE, and quantifying the DNA pellet by Qubit HS fluorimetry. Ogataea cells from a fresh pellet containing ~3.5µg DNA were resuspended in 270µL 1x *Dpn*II digestion buffer (50mM Tris HCl pH 7,9, 100mM NaCl, 10mM MgCl2, 100µg/mL Bovine Serum Albumin [BSA]) and vortexted with hydrated glass beads (Sigma Aldrich cat# G1145) for 5 minutes (30 seconds vortexing, 30 seconds on ice) to lyse cells. Then, 270uL of supernatant and cell debris was moved to a new tube, SDS was added to 0.625%, samples were incubated at 62°C for 7 minutes, and Triton X-100 was added to 1%. Chromatin was digested with 200U *Dpn*II (NEB cat# R0543L) for overnight at 37°C. DNA was end-repaired with 25U Klenow (NEB cat# M0210L) with 30uM dTTP, dCTP, dGTP, and 28uM biotin-14-dATP (Invitrogen cat# 19524-016) in 1x *Dpn*II buffer and ligated with T4 DNA ligase (1675U; NEB cat# M0202L) in 1x Ligation buffer with 1% Triton X-100 and 100 µg/mL BSA. Ligated DNA fragments were treated once with 600ug/mL proteinase K overnight at 65°C and again for 2 hours at 65°C the following morning, organically extracted with phenol/chloroform/IAA, ethanol precipitated with 40µg [final] glycogen at -80°C, resuspended in 50µL TE, and treated with 400ug/mL RNaseA at 37°C for 30 minutes. DNA ligation products were organically extracted with phenol/chloroform/IAA (25:24:1) and chloroform, and ethanol precipitated at -80°C, and resuspended in 25µL TE/10 (10mM Tris-HCl pH 8.0; 0.1mM EDTA). Unligated biotinylated nucleotides were removed with

5U T4 DNA polymerase (NEB cat# M0203L) in the presence of 100uM dATP and dGTP in 1x NEB Buffer 2.1 at 12°C for 2 hours, quenched with 10mM EDTA, organically extracted with phenol/chloroform/IAA (25:24:1) and chloroform, ethanol precipitated at -80°C, resuspended in 500µL TE/10, and sheared by sonication with a QSonica sonicator (model# Q55) for five minutes, using a 30 seconds sonication, 30 seconds on ice cycle at amplitude 30. Biotinylated DNA ligation fragments were purified with 25µL Streptavidin M280 Dynabeads (ThermoFisherScientific cat# 11205D) resuspended in 50µL 2x BW buffer (10mM Tris-HCL pH 8.0, 1mM EDTA, 2M NaCl) with an additional 450µL 2x BW buffer for 30 minutes at room temperature. Streptavidin beads were washed twice with 1mL 1x BW buffer, once with 200µL 1x BW buffer, and twice with 200µL TE/10; beads were resuspended in 25µL TE/10.
