## Supplemental_Table_S1 for "A Translocation within the *Ogataea* Species Complex Alters Local Subtelomeric Chromatin while Maintaining Overall Genome Organization"

Table S1. Class III NAD+ Dependent Deacetylases in *Ogataea* Genomes

| | | *O. polymorpha* | | *O. haglerorum* | |
|---|---|---|---|---|---|
| HDAC | *S. cerevisiae* RefSeq | Homolog RefSeq | blastp e value vs. *S. cerevisiae* protein | Homolog | blastp e value vs. *S. cerevisiae* protein |
| *SIR1* | NP_013027 | - | | - | |
| *SIR2* | NP_010242 | - | | - | |
| *SIR3* | NP_013547 | - | | - | |
| *SIR4* | NP_010513 | - | | - | |
| *ORC1* | NP_013646 | XP_018212351 | 5E-94 | KL911_001080 (complement(2:788349..790556)) | 2E-95 |
| *HST1* | NP_014573 | XP_018209953 | 4E-144 | KL911_001176 (5:648227..649822) | 4E-141 |
| *HST2* | NP_015310 | XP_018211510 | 3E-104 | KL911_000064 (complement(3:785654..786640)) | 2E-100 |
| *HST3* | NP_014668 | XP_018208924 | 2E-115 | KL911_004240 (7:758477..759649) | 4E-112 |
| *HST4* | NP_010477 | XP_018211420 | 2E-103 | KL911_000162 (3:635067..636218) | 1E-98 |
