## Supplemental_Table_S2 for "A Translocation within the *Ogataea* Species Complex Alters Local Subtelomeric Chromatin while Maintaining Overall Genome Organization"

Table S2: Genome Statistics for the *O. polymorpha* and *O. haglerlorum* genomes

| | *Ogataea polymorpha* | *Ogataea haglerorum* |
|---|---|---|
| NCBI RefSeq assembly | GCF_001664045.1 | n/a |
| total genome size | 8,974,850 bp | 8,739,724 bp |
| Total ungapped length | 8,974,850 bp | 8,739,724 bp |
| number of scaffolds | 7 | 7 |
| number of contigs | 7 | 7 |
| Scaffold N50 | 1,302,532 bp | 1,285,713 bp |
| Scaffold L50 | 4 | 4 |
| Number of contigs | 7 | 7 |
| GC percent | 47.86 | 49.65 |
| # N's per 100 kb | 5.37 | 19.45 |
